# Cryptic variation fuels plant phenotypic change through hierarchical epistasis

**DOI:** 10.1101/2025.02.23.639722

**Authors:** Sophia G. Zebell, Carlos Martí-Gómez, Blaine Fitzgerald, Camila Pinto Da Cunha, Michael Lach, Brooke M. Seman, Anat Hendelman, Simon Sretenovic, Yiping Qi, Madelaine Bartlett, Yuval Eshed, David M. McCandlish, Zachary B. Lippman

## Abstract

Cryptic genetic variants exert minimal or no phenotypic effects alone but have long been hypothesized to form a vast, hidden reservoir of genetic diversity that drives trait evolvability through epistatic interactions. This classical theory has been reinvigorated by pan-genome sequencing, which has revealed pervasive variation within gene families and regulatory networks, including extensive cis-regulatory changes, gene duplication, and divergence between paralogs. Nevertheless, empirical testing of cryptic variation’s capacity to fuel phenotypic diversification has been hindered by intractable genetics, limited allelic diversity, and inadequate phenotypic resolution. Here, guided by natural and engineered cis-regulatory cryptic variants in a recently evolved paralogous gene pair, we identified an additional pair of redundant trans regulators, establishing a regulatory network that controls tomato inflorescence architecture. By combining coding mutations with a cis-regulatory allelic series in populations segregating for all four network genes, we systematically constructed a collection of 216 genotypes spanning the full spectrum of inflorescence complexity and quantified branching in over 27,000 inflorescences. Analysis of the resulting high-resolution genotype-phenotype map revealed a layer of dose-dependent interactions within paralog pairs that enhances branching, culminating in strong, synergistic effects. However, we also uncovered an unexpected layer of antagonism between paralog pairs, where accumulating mutations in one pair progressively diminished the effects of mutations in the other. Our results demonstrate how gene regulatory network architecture and complex dosage effects from paralog diversification converge to shape phenotypic space under a hierarchical model of epistatic interactions. Given the prevalence of paralog evolution in genomes, we propose that paralogous cryptic variation within regulatory networks elicits hierarchies of epistatic interactions, catalyzing bursts of phenotypic change.

**Keyword:** cryptic mutations, paralogs, redundancy, cis-regulatory, tomato, inflorescence, gene regulatory network, modeling, epistasis

## INTRODUCTION

An enduring debate in evolutionary biology concerns the extent to which small-effect genetic variants contribute to expanding phenotypic diversity from developmentally stabilized (canalized) states ^1–4^. The most enigmatic cohort among small-effect variants may be cryptic alleles ^5,6^. In their simplest form, cryptic alleles only have substantial effects on phenotypes through epistatic interactions with other alleles, including other cryptic alleles ^7^. Cryptic variation is most likely to accumulate in buffered molecular contexts, such as redundancy within gene families as well as within gene regulatory networks ^8^. The accumulation of cryptic alleles in these contexts may be a major source of genomic variation shaping the trajectories of phenotypic evolution. Under this hypothesis, cryptic variant epistasis may mediate adaptive potential and, in some cases, elicit novel morphologies, facilitating both within-species adaptation and macroevolutionary transitions ^8–10^.

Demonstrating cryptic variation’s potentially widespread contribution to trait evolution is challenging. Genetic dissection of trait variation is typically confined to single species or a few closely related ones, where only narrow ranges of phenotypic diversity can be assessed. Moreover, most dissections of natural trait variation expose only major effect variants, as is true for developmental genetics in model systems, leaving the influence of cryptic alleles on natural populations and gene regulatory networks largely unexplored ^6,11^. Importantly, background dependencies–likely stemming in part from cryptic alleles–are common in evolutionary and developmental genetics ^12–15^. Yet, despite these clues, efforts to systematically dissect cryptic variation and its potential to drive phenotypic change remain hindered by the intricate and often poorly understood structure and redundancy of gene regulatory networks, combined with limited allelic diversity and constrained phenotypic resolution in most systems.

Genome editing in model systems with complex developmental programs offers a powerful approach to interrogate cryptic variation. Beyond applications in medicine and agriculture, genome editing enables the precise engineering of customized mutations and allelic series with a wide range of phenotypic effects in controlled isogenic backgrounds ^16^. This capability allows for deeper exploration of gene function and interactions among different types of mutations, including *cis*-regulatory variants and paralog duplication and loss that influence dosage but are often cryptic ^17–20^. While pairwise interactions are often detectable, how diverse allelic variants interact across larger regulatory and developmental networks remains unexplored.

Natural variation in the architectures of plant reproductive branching systems (inflorescences) within the Solanaceae family, particularly in the *Solanum* genus, exemplifies how evolutionary processes generate morphological diversity (**Fig. 1a**)^21,22^. Cryptic variation may shape such trait diversity, making tomato (*Solanum lycopersicum*) an ideal system to test this hypothesis within a controlled genetic framework. Many tomato mutants—often involving cryptic genetic interactions—mirror natural variation across the family, offering a platform to systematically dissect how cryptic variation can influence inflorescence architecture ^23^. We uncovered epistasis between cryptic natural coding and cis-regulatory variants in two MADS-box transcription factor genes of the *SEPALLATA* (*SEP*) clade, which have conserved roles in specifying flower identity (**Fig. 1b**) ^24,25^. Interactions between mutations in the *SEP* gene *JOINTLESS2* (*J2*), originating from the wild species *S. cheesmaniae*, and a natural *cis-*regulatory variant in its paralog *ENHANCER OF JOINTLESS2* (*EJ2*), result in highly branched inflorescences through classical redundancy epistasis ^24^. While individual mutations in each paralog are cryptic on inflorescence branching, different combinations of homozygous and heterozygous genotypes produce varying degrees of branching effects, reflecting a dose-dependent epistatic relationship (**Fig. 1b**). Notably, while *EJ2* is conserved across the Solanaceae, *J2* is absent in many species and cultivated genotypes, making them sensitive to changes in inflorescence architecture from variations in *EJ2* dosage ^26^. The *J2*-*EJ2* paralog relationship offers a system to study how cryptic variation in a broader regulatory network influences epistasis and trait evolvability. However, realizing this potential first requires identifying additional network components.

**Fig. 1:**
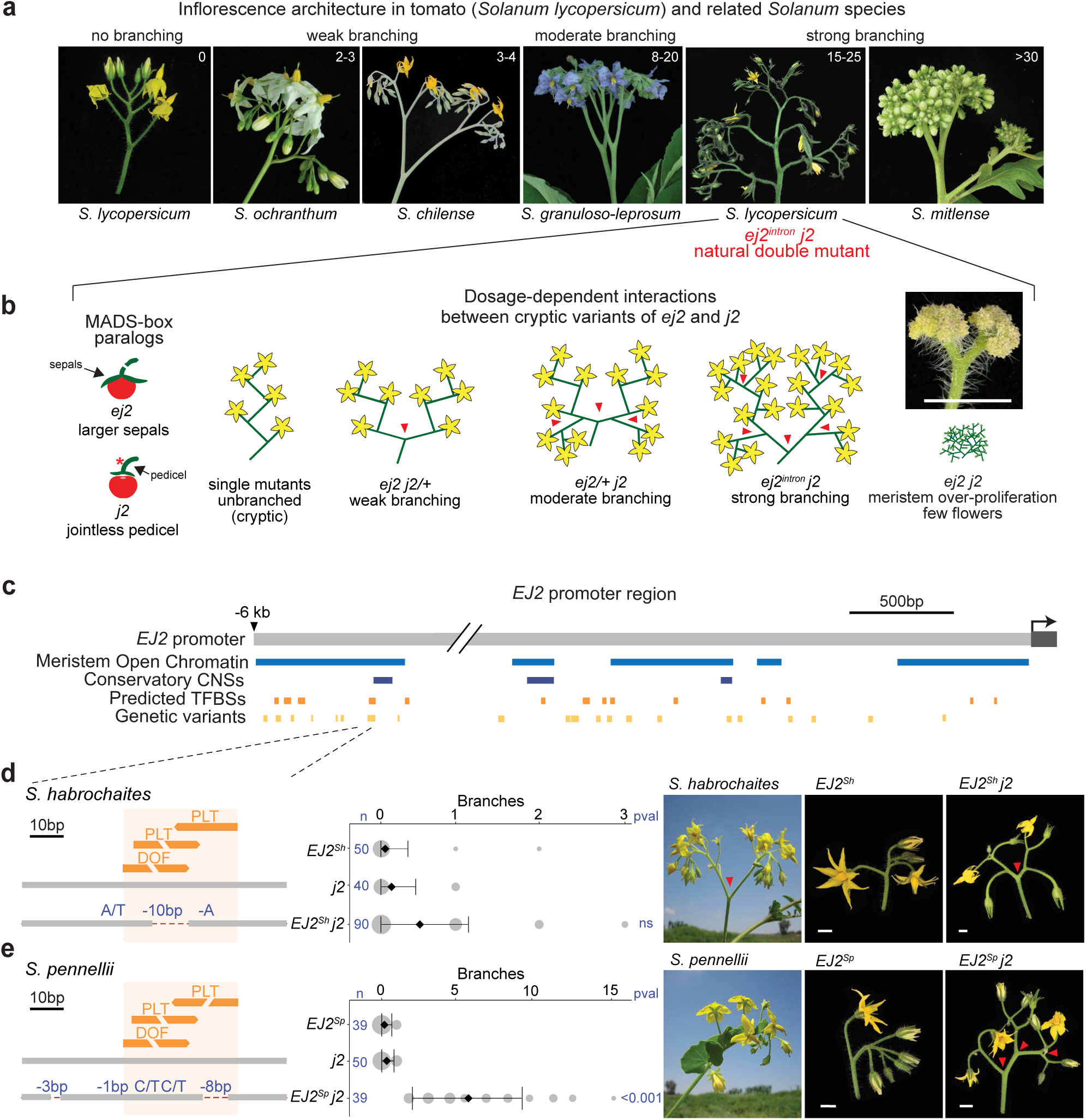
Natural *cis*-regulatory variants of the *SEP* gene *EJ2.* **(a)** Increased complexity of inflorescences of *Solanum* species. **(b)** Dose-dependent redundancy relationship among the SEP paralogs *EJ2* and *J2* in controlling tomato inflorescence architecture. **(c)** A 6 kb region upstream of the *EJ2* transcription start site showing open chromatin (blue), conserved non-coding sequences (CNSs, dark blue), predicted transcription factor binding sites (TFBSs, orange), and pan-genome variants (light orange). **(d)** *Cis*-regulatory region of *S. lycopersicum* and *S. habrochaites EJ2* with one DOF and two PLT TFBSs. The overlapping DOF-PLT site is disrupted by a 10 bp deletion in *S. habrochaites*. Quantification of inflorescence branching for the three indicated genotypes (middle) is followed by representative images of *S. habrochaites*, the *EJ2-*containing introgression line (*EJ2^Sh^*), and *EJ2^Sh^* in the *j2* background (*EJ2^Sh^ j2,* right). **(e)** *Cis*-regulatory region of *S. pennellii* showing disruption of all three TFBSs by a 10 bp deletion and linked SNVs, quantification of inflorescence branching for the three indicated genotypes (middle) and representative images of *S. pennelli*, the *EJ2-*containing introgression line (*EJ2^Sp^*), and *EJ2^Sp^* in the *j2* background (*EJ2^Sp^ j2,* right). For ‘d’ and ‘e’: Dashed red lines, deleted sequences; Blue text, deletion sizes and SNVs. Area of grey circles, numbers of inflorescences quantified; black diamonds and bars, mean values and standard deviations. Total number of inflorescences (n) and p-values from Two-sided Dunnett’s Compare with Control Test. Red arrowheads mark branch points. Scale bars are 1 cm.

## RESULTS

### Cryptic variants of the MADS-box gene *EJ2*

To examine whether natural variation in the epistatic relationship between *J2* and *EJ2* could further contribute to inflorescence architecture diversity, we mined tomato pan-genome data for variation in the *EJ2* promoter ^27,28^. Using open chromatin in tomato reproductive meristems and its overlap with transcription factor binding sites (TFBSs) (**Fig. 1c**) ^29^, we filtered 629 candidate variants (**Supplementary Table 1**) down to two small deletions and nearby single-nucleotide variants (SNVs) that coincided with a cluster of three TFBSs located 6 kbp upstream of *EJ2* (**Fig. 1c,d, Supplementary Table 2**). These variants were found only in the wild species *S. habrochaites* and *S. pennellii*, which produce weakly branched inflorescences (**Fig. 1d,e**).

Tomato introgression lines carrying wild species chromosomal segments with these *EJ2* variants in isogenic backgrounds (designated *EJ2^Sh^* and *EJ2^Sp^*) rarely exhibit branching (**Fig. 1d,e**). However, branching increased with the addition of the *j2* mutation. While *EJ2^Sh^ j2* plants exhibited a subtle, though not significant, increase in branching compared to the *EJ2^Sh^* genotype (**Fig. 1d**), *EJ2^Sp^ j2* plants produced an average of five branches per inflorescence (**Fig. 1e, Supplementary Tables 3-4**). Since introgression lines have large chromosomal segments carrying additional variants, we used CRISPR to test whether branching specifically resulted from TFBS disruption by attempting to create similar deletions in *j2* mutant background. Due to the absence of Cas9 gRNA target sites in the 25 bp region, we first used the more permissive Cas9 SpRY variant, which recognizes an expanded protospacer-adjacent motif (PAM) ^30,31^. This approach, which used three gRNAs and catalytically active and dead versions of Cas9 fused to an adenine base editor, produced three alleles, each with a single SNV within one TFBS. As none of these single nucleotide changes led to branching (**Supplementary Fig. 1, Supplementary Tables 5, 6**), we targeted a 153 bp target region flanking the TFBSs using four conventional Cas9 gRNAs. We recovered seven *EJ2* promoter (*EJ2^pro^*) alleles (**Fig. 2a**): five with small indels and SNVs, often at the gRNA target sites, and two with overlapping ∼100 bp deletions spanning the entire interval. Notably, none of these alleles exhibited branching in the functional *J2* background, but all caused branching in the mutant *j2* background, displaying a continuous range of effects (**Fig. 2b, Supplementary Tables 7, 8)**. Additionally, none of these genotypes exhibited the pleiotropic phenotypes observed in loss of function double mutants, such as enlarged sepals or altered fruit shape (**Supplementary Fig. 2, Supplementary Tables 9-12**) ^24^. These findings suggest that these TFBSs along with nearby sites affected in several alleles collectively act as an enhancer regulating *EJ2* expression to control meristem maturation and inflorescence development.

**Fig. 2:**
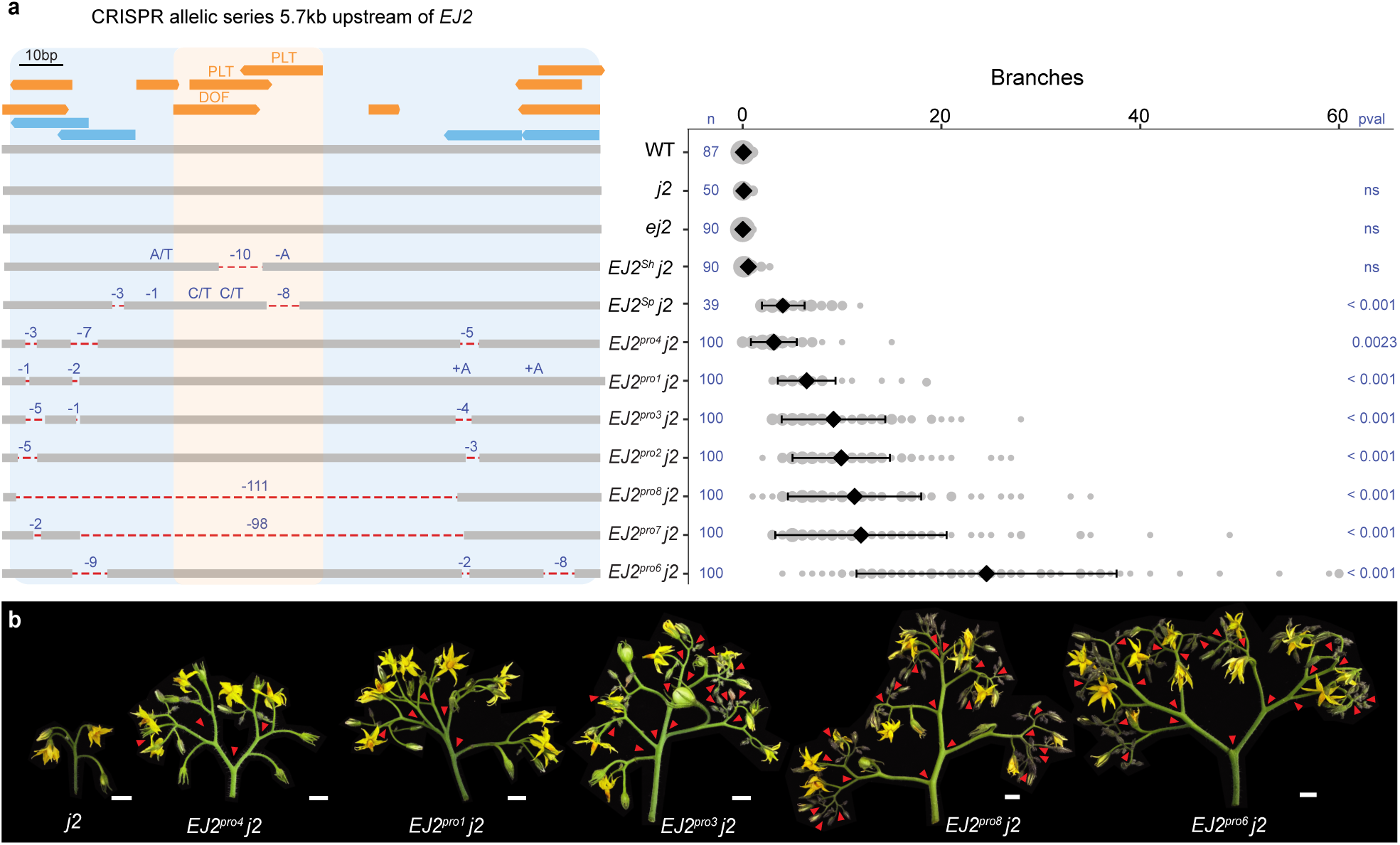
Engineered *EJ2 cis*-regulatory alleles pinpoint a branching specific enhancer. **(a)** Left: 153 bp target region located 5.7kb upstream of the *EJ2* translation start site showing annotated TFBSs (orange), including the focal DOF and PLT binding sites, and four CRISPR/Cas9 gRNAs (light blue) used for generation of the *EJ2* promoter (*EJ2^pro^*) allelic series. The lines with the strongest phenotypes (*EJ2^pro7^ j2* and *EJ2^pro8^ j2*) carried overlapping ∼100 bp deletions that removed the DOF-PLT binding sites, as well as other lower confidence predicted binding sites (**Supplementary Table 2, 22**). Red lines, deleted sequences; Deletion sizes and SNVs, blue. Right: Quantifications of inflorescence branching for each genotype. Area of grey circles, numbers of inflorescences; black diamonds and bars, mean values and standard deviations. Total number (n) of observed inflorescences and p-values from Two-sided Dunnett’s Compare with Control Test. **(b)** Representative images of *j2* and five of the *EJ2^pro^* alleles in the *j2* background, capturing the range of branching effects. Red arrowheads mark branch points. Scale bars are 1 cm.

### *PLT* paralogs regulate *EJ2* and branching

Our finding that multiple *cis*-regulatory cryptic alleles caused branching, partly due to the disruption of the TFBSs affected by two natural alleles, prompted us to investigate whether the transcription factors predicted to bind these specific sites directly regulate *EJ2* expression and inflorescence architecture. Both the *S. habrochaites* and *S. pennellii EJ2 cis*-regulatory alleles are predicted to disrupt binding sites for the DOMAIN OF UNKNOWN FUNCTION (DOF) and the PLETHORA (PLT) transcription factor families. Members of these families have been implicated in meristem development in *Arabidopsis thaliana* (PLT) ^32^ and flowering in tomato (DOF) ^33^. Using our tomato meristem maturation expression atlas ^34^, we searched for *DOF* and *PLT* genes expressed during the transition meristem stage. Among the 34 *DOF* genes in tomato*, SlDOF9* emerged as a primary candidate (**Supplementary Fig. 3**), supported by prior findings that engineered mutants of this gene develop more flowers on inflorescences with weak branching. However, our CRISPR mutants exhibited a drastic change in leaf shape but did not show branching, either alone or in the *j2* background (**Supplementary Fig. 3, Supplementary Tables 13,14**).

We next focused on two closely related *PLT* paralogs (*SlPLT3* and *SlPLT7;* hereafter *PLT3* and *PLT7*), expressed in meristems during and after floral transition, similar to *J2* and *EJ2* (**Fig. 3a**). These tomato *PLT* paralogs are orthologs of Arabidopsis *AtPLT3* and *AtPLT7* but arose from independent duplications (**Fig. 3b**). In Arabidopsis, *AtPLT3* and *AtPLT7* function in meristem maturation as well as floral organ identity and growth (Horstman 2014). We tested whether the PLT proteins bind the *EJ2* promoter and activate its expression by performing a heterologous dual-luciferase assay in tobacco (*Nicotiana benthamiana*) leaves. Although the full-length coding sequence of *PLT7* could not be cloned or synthesized, PLT3 strongly activated the intact *EJ2* promoter but not the *EJ2^pro8^*, *EJ2^pro-Sh^* and *EJ2^pro-Sp^* alleles, which have mutated PLT binding sites (**Fig. 3c, Supplementary Tables 15, 16**). For normalization, we included ETHYLENE RESPONSIVE12 (ERF12) as a non-binding control, a transcription factor expressed in meristems with a single DNA-binding domain structurally similar to those in the PLTs.

**Fig. 3:**
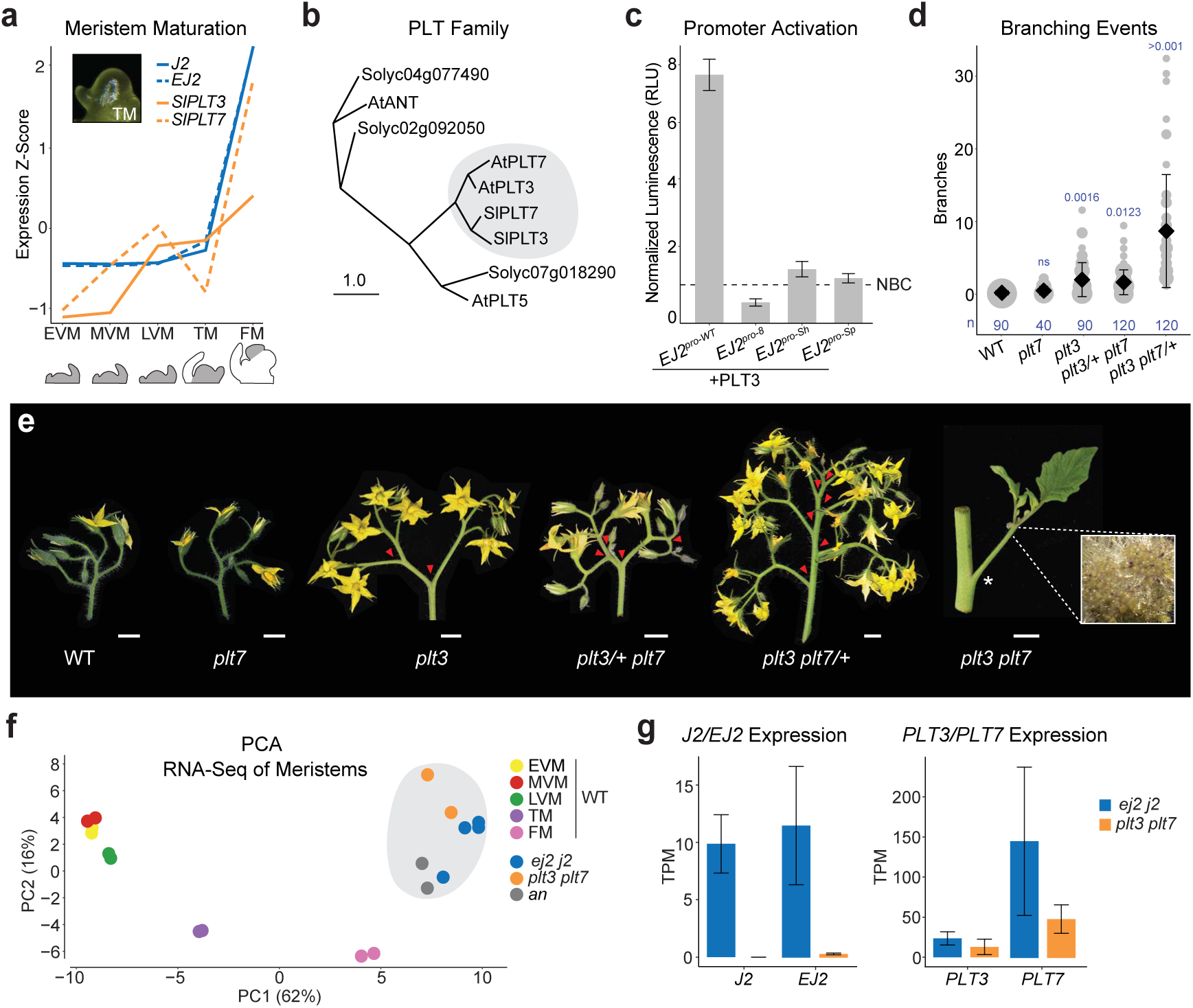
*PLETHORA* transcription factor paralogs control inflorescence architecture upstream of *EJ2/J2*. **(a)** Meristem expression dynamics of *EJ2, J2, PLT3,* and *PLT7* over five depicted (grey) stages: Early Vegetative Meristem (EVM), Middle (MVM), Late (LVM), Transition (TM), and Floral Meristem (FM). **(b)** Phylogenetic tree of PLT proteins from Arabidopsis (*At*) and tomato (*Sl*). The paralog pairs AtPLT3/SlPLT3 and AtPLT7/SlPLT7 are circled. Scale bar is relative substitution rate. **(c)** *In planta* heterologous expression of firefly luciferase driven by wild type and *EJ2 cis*-regulatory alleles, co-transfected with PLT3 and normalized to an internal constitutive renilla luciferase and nonbinding protein controls (NBC). **(d)** Quantification of branching in double *plt3/7* mutants, including single and homozygous/heterozygous combinations. Grey circles correspond to the number of inflorescences, black diamonds are mean values and bars are standard deviations. Number (n) of observations and p-values from Two-sided Dunnett’s Compare with Control Test shown in blue. **(e)** Representative inflorescences of *plt3* and *plt7* mutant combinations. Inset is double mutant inflorescence showing meristem over-proliferation (5x magnification). Red arrowheads mark branch points. Scale bars are 1 cm. **(f)** Principal Component Analysis of top 200 genes differentially expressed over meristem maturation in staged WT meristems and proliferating meristems of the mutant genotypes: *ej2 j2, plt3 plt7,* and *an.* **(g)** Expression of paralogs *EJ2, J2, PLT3,* and *PLT7* in proliferating meristems of the mutants *ej2 j2* and *plt3 plt7* in transcripts per million.

We mutated both *PLT* paralogs using CRISPR/Cas9. *plt7* single mutants appeared wild type, while *plt3* mutants rarely produced branched inflorescences. However, double mutants exhibited extreme meristem overproliferation and branching. These mutant alleles also displayed dose-dependent redundancy: *plt3/+ plt7* genotypes showed weak branching, while *plt3 plt7/+* genotypes exhibited moderate branching (**Fig. 3d,e, Supplementary Table 17, 18**). The binding assays, combined with the quantitative effects of the *plt* mutant genotypes and the intermediate branching observed in all *EJ2pro j2* genotypes, suggest that the PLTs transcriptionally regulate *EJ2* and likely other genes involved in inflorescence development. This aligns with the presence of PLT binding sites in the *cis*-regulatory regions of *J2* (**Supplementary Table 19**).

To further characterize the functional relationships between the *PLT* and *SEP* paralogs, we profiled and compared gene expression in proliferating meristems from the two double mutants. A principal component analysis (PCA) of the top 200 maturation marker genes from wild type meristem stages ^34^ revealed that both double mutants cluster closest to the floral meristem maturation stage and also with the *anantha* (*an*) mutant, which overproliferates floral meristems due to a mutation in the ortholog of the Arabidopsis *UNUSUAL FLORAL ORGANS* gene (**Fig. 3f**). The expression data also showed that *J2* and *EJ2* are not expressed in *plt3 plt7* meristems, whereas both *PLTs* remain expressed in *j2 ej2* meristems, supporting regulation of these *SEPs* by the PLTs (**Fig. 3f,g)**.

### A *PLT-SEP* genotype-phenotype map

Our genetic and molecular analyses identified key components of an inflorescence regulatory network comprising two duplicated transcription factor pairs with dose-dependent epistatic interactions. We generated multiple alleles for these factors in a shared isogenic background, including both promoter regulatory site alleles and strong coding alleles, all of which are cryptic individually. This genetic resource enabled us to systematically explore the phenotypic space and quantify the functional output arising from variations within a simple, rapidly evolving network involving the interactions among *PLT3* and *PLT7*, and their downstream targets, *J2* and *EJ2*. (**Fig. 4a**).

**Fig. 4:**
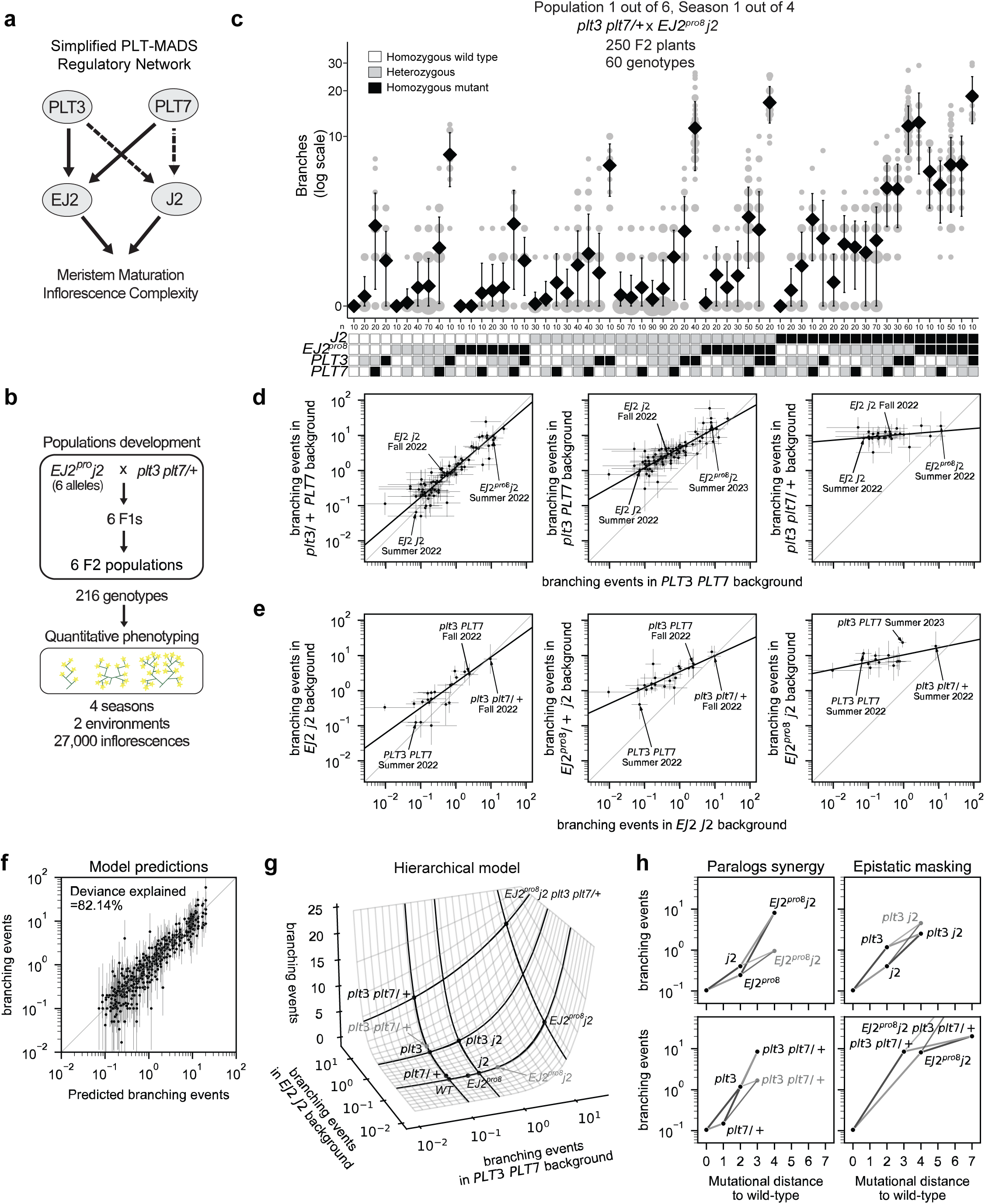
A hierarchical model of genetic interactions best explains the *PLT-SEP* inflorescence genotype-phenotype map. **(a)** Simplified architecture of the *PLT3/7-EJ2/J2* genetic and molecular network. Arrows represent positive regulation. Solid arrows are supported by genetic and molecular evidence. Dashed arrows are supported by annotated binding sites and molecular evidence. **(b)** Six F2 populations with different *EJ2^pro^* alleles were phenotyped across multiple seasons and environments, yielding observations of per-inflorescence branching events for 216 different genotypes. **(c)** Representative data for inflorescence branching from a single season in one of six populations. The number of branching events is represented in a log(1+x) scale, with tick labels giving the original x values. Area of grey circles correspond to the number of inflorescences, black diamonds and bars denote mean values and standard deviations. Total number (n) of observations. Genotypes are represented in a matrix below, showing all 60 homozygous (black) and heterozygous (gray) mutant combinations with observed counts data. **(d, e)** Maximum Likelihood Estimates (MLEs) of the mean number of branching events. Each dot represents a combination of mutations in *EJ2* and *J2* (d) or *PLT3* and *PLT7* (e) in a specific season. The plots show the combinations of mutations across different genetic backgrounds of mutations in *PLT3* and *PLT7* (d) or *EJ2^pro8^* and *J2* loci (e) with the x-axis always showing the corresponding phenotype on a wild-type background. Error bars represent the 95% confidence interval for each genotype-season combination. Genotype-season combinations with a 95% confidence interval wider than a thousand-fold range are not shown. Black dashed lines represent the total least squares regression lines of the estimated log-branching events. **(f)** Comparison of the predicted number of branching events by the hierarchical model and the MLE for each genotype-season combination. Error bars represent the 95% confidence interval for the MLEs. Genotype-season combinations with a 95% confidence interval wider than a thousand-fold range are not shown. **(g)** Representation of the inferred hierarchical epistasis model. The surface represents how the phenotypes conferred by perturbing each pair of paralogs in the wild-type background of the other pair are combined together to determine the average phenotype for genotypes with mutations across both pairs of paralogs. Transects along the surface represent different genetic backgrounds (WT, *j2* and *EJ2^pro8^j2* on one axis and WT, *plt3* and *plt3 plt7/+* on the other), which become shallower in increasingly mutated backgrounds, illustrating the phenomena of epistatic masking. Black points in the surface represent predictions of the hierarchical model for the labeled genotypes, whereas grey dots represent the multiplicative expectation for double mutants in each pair of paralogs. **(h)** Predicted branching events under the hierarchical model in specific genotypes as a function of their mutational distance to the wild-type. Grey dots and dashed lines represent the multiplicative expectation for combining the individual sets of mutations.

To explore the genotype-phenotype map of this tomato inflorescence regulatory network, including the interplay between cryptic dosage effects and paralogous epistatic relationships, we selected six *EJ2^pro^ j2* lines spanning a range of branching effects and crossed them with *plt3 plt7/+* plants to generate six F2 segregating populations (**Fig. 4b, Methods**). These populations provided 216 genotypic combinations, enabling an in-depth phenotypic and statistical analysis of branching effects from single, double, and higher-order mutants, as well as dosage effects from heterozygosity (**Fig. 4c**). Across four field seasons in two environments, we quantified over 27,000 inflorescences (**Supplementary Table 20**). Preliminary analyses indicated greater variance in the number of branching events per inflorescence both within plants and within genotypes than would be expected if branching events were Poisson distributed (**Supplementary Fig. 4a,b)**. Consequently, we treated branching events as overdispersed count data in all subsequent analyses, which provided a significantly improved fit relative to a Poisson error model (likelihood ratio test for negative binomial versus Poisson: p < 10^-16^; Methods). For illustration, data from a single population and field season are shown in **Fig. 4c**, with the full dataset available in **Supplementary Table 20**.

Using the data from this large set of crosses, we then sought to determine how mutations within this genetic network combine to determine the mean number of branching events per inflorescence for any given genotype. We began by comparing a model where mutations at different loci combine additively versus a model where mutations at different loci combine multiplicatively (with both models accounting for dominance interactions within each locus). The model with multiplicative effects fit substantially better (59.40% deviance explained by an additive model with a log link versus 46.05% deviance explained by an additive model with an identity link **Supplementary Fig. 4c,d**), suggesting an overall tendency for mutations at different loci to interact multiplicatively in this system. Using this multiplicative model as a baseline, we then fit a more complex model to determine whether epistatic interactions between loci are also present. We found that a pairwise interaction model that included these epistatic interactions significantly outperformed the multiplicative model (83.11% deviance explained, likelihood ratio test: p < 10^-16^) and achieved greater predictive performance for held-out seasons and genotypes (**Supplementary Fig. 5**; average leave-one-out cross-validated R^2^ on held-out seasons was 0.89 for the pairwise model compared to 0.70 for the multiplicative model). This pairwise model detected pervasive epistasis across loci, with 43 of 80 epistatic terms showing *p* values < 0.05 (**Supplementary Table 21**). Among the most notable of these interactions were strong, super-multiplicative, synergistic interactions between *plt3* and *plt7* as well as between the *EJ2^pro^* alleles and *j2*, such that combining mutations within a paralog pair often results in a mean number of branching events several fold (3- to 9) in excess of what would be expected when multiplying the effects of the individual mutations (**Table 1**).

**Table 1.** Estimates of fold change relative to the expectation under the multiplicative model for combinations of perturbations within each paralog pair under the pairwise interaction model. For the full set of estimated parameters, see Supplementary Table 21.

| Genotype | Fold Change in Excess of Multiplicative Expectation | 95% Confidence interval | P-value |
| --- | --- | --- | --- |
| *plt3/+ plt7* | 3.29 | [2.64, 4.10] | $1.44 \cdot 10^{-26}$ |
| *plt3 plt7/+* | 3.27 | [2.65, 4.04] | $2.95 \cdot 10^{-28}$ |
| *EJ2<sup>pro3</sup> j2* | 6.81 | [4.21, 11.03] | $5.16 \cdot 10^{-15}$ |
| *EJ2<sup>pro4</sup> j2* | 9.32 | [5.25, 16.57] | $2.74 \cdot 10^{-14}$ |
| *EJ2<sup>pro1</sup> j2* | 7.05 | [5.21, 9.55] | $9.91 \cdot 10^{-37}$ |
| *EJ2<sup>pro7</sup> j2* | 16.97 | [8.14, 35.38] | $4.16 \cdot 10^{-14}$ |
| *EJ2<sup>pro8</sup> j2* | 5.95 | [4.42, 8.03] | $1.13 \cdot 10^{-31}$ |
| *EJ2<sup>pro6</sup> j2* | 6.49 | [4.56, 9.23] | $3.10 \cdot 10^{-25}$ |

While we observed strong positive interactions within paralog pairs, consistent with our previous observation regarding at least partial redundancy, we also inferred many negative interactions between non-paralogous pairs (11 out of 14 additive-by-additive epistatic terms between non-paralogous pairs were negative and showed *p* values < 0.05, **Supplementary Table 21**). To better understand the structure of the genotype-phenotype relationship implied by these negative interactions, we first calculated maximum likelihood estimates for the average number of branching events per inflorescence across the 470 genotype-season combinations in our data set. We then plotted how the phenotypes conferred by different combinations of *j2* and *EJ2^pro^* mutations are transformed when placed on different *plt3* and *plt7* mutant backgrounds as compared to a wild-type *PLT3* and *PLT7* background. Interestingly, after log-transformation, we observed that the relationship between backgrounds is always linear; however, the slope of this linear relationship decreased as mutations accumulated in the *PLT3* and *PLT7* loci (**Fig. 4d, Extended Data Fig. 1**). We observed the same pattern when analyzing the phenotypes resulting from *plt3* and *plt7* mutant combinations across different *j2* and *EJ2^pro8^* backgrounds, with slopes decreasing as the strength of the *EJ2 J2* perturbations increased (**Fig. 4e**). Importantly, this pattern was consistent across all six *EJ2^pro^* alleles (**Extended Data Fig. 1**) and remained robust across other methods of estimating average phenotype (**Extended Data Fig. 2**). Collectively, these analyses suggest a surprisingly simple form of genetic interaction between the paralog pairs, where mutations in one paralog pair linearly re-scale the effects of mutations in the other. As the slope of the linear relationship reduces almost to zero in highly mutated genetic backgrounds, the negative interaction coefficients between the paralog pairs reflect a systematic pattern of masking interactions between non-paralogous mutations, wherein the effects of mutations in one pair is diminished when the other pair is highly mutated.

### Hierarchical epistasis in a genotype-phenotype map

This mode of genetic interaction, in which each mutation systematically re-scales the effects of other mutations, is a defining feature of a classic theoretical model of epistasis known as the multilinear model ^35^. To simultaneously capture the super-multiplicative synergistic interactions within paralog pairs together with the systematic masking interactions between paralog pairs, we thus fit an additional model that we call the hierarchical epistasis model because it treats this multilinear interaction between paralog pairs as an additional layer of epistasis. In this model, similar to the pairwise model, we allowed an arbitrary pattern of dominance and epistasis within each paralog pair, but the predictions based on the coefficients for each pair of paralogs are combined according to a multilinear interaction. Crucially, because for a multilinear interaction mutations in one of the pairs of paralogs simply rescale the effects of mutations in the other pair, this multilinear interaction between paralog pairs uses only a single parameter to describe the extent to which mutations in one paralog pair mask the effects of mutations in the other pair, in contrast to our more traditional pairwise interaction model that uses 42 parameters to describe interactions between the paralog pairs (i.e. additive-by-additive, dominance-by-dominance, and additive-by-dominance parameters for each of the 14 non-paralogous pairs of mutations). Finally, after the within-paralog pair interactions are combined across paralog pairs using a multilinear interaction, the results are transformed once more using an exponential function (i.e. we again use a log link, identical to that used for the pairwise model, see **Methods**). We applied this hierarchical epistasis model to our data and found it recapitulated the observed phenotypic measurements nearly as well as the full pairwise model (82.14% deviance explained, **Fig. 4f**), and maintained high predictive power for held-out data from complete seasons and unobserved genotypes (**Supplementary Fig. 5**, average leave-one-season-out cross-validated R^2^=0.89), despite the substantially reduced number of parameters. Using the Akaike Information Criterion (AIC) to compare the two models ^36,37^, we found that the hierarchical model is roughly 18,358 times more likely than the pairwise model (AIC_hiearchical_ - AIC_pairwise_ = -19.64).

To provide a more intuitive understanding of the behavior of this hierarchical model, it is useful to visualize the combination of interaction between paralog pairs and the exponential mapping as a response surface ^38^ that depicts the predicted phenotype of a combination of mutations across both paralog pairs as a function of the phenotypes conferred when mutating each paralog pair separately (**Fig. 4g, Extended Data Fig. 3a**). In this response surface, the phenotypes conferred by within-pair mutations are shown on a log scale, so that the effects of mutations that combine according to the multiplicative expectation would add along each axis. However, consistent with the pairwise model, interactions within each paralog pair show a strong pattern of synergy (**Fig. 4h, left,** see **Extended Data Fig. 3b** for all other within paralog pair interactions), such that double mutants are substantially further displaced along each axis than would be expected from the individual effects of the single mutants (**Fig. 4g**, multiplicative expectation shown by gray dots). The key difference from the pairwise model is that the interactions between paralog pairs, instead of being represented by numerous between-pair interaction terms, are given by a surface (**Fig. 4g**) whose shape is controlled by a single parameter that determines the overall strength of the interaction between the paralog pairs, and which has horizontal and vertical transects that are exponential due to the final exponentiation (**Extended Data Fig. 3b**). In particular, we see that as the phenotypic effect of one paralog pair increases, the corresponding transect for mutations in the other pair becomes progressively flatter, reflecting diminishing phenotypic contributions from additional mutations (**Extended Data Fig. 3c)**. While this masking effect is modest for mutations with moderate phenotypic impacts (e.g., the 11.24-fold effect of a homozygous *plt3* mutation in a wild-type background is reduced to 6.18-fold in a *j2* background, **Fig. 4h, top right**), it becomes far more pronounced in highly mutated backgrounds (e.g., adding *plt3 plt7/+* to a wild-type *EJ2/J2* background has an 81.30-fold effect, but only a 2.49-fold effect in the *EJ2^pro8^ j2* background, **Fig. 4h, bottom right**). Overall, these results show how synergy within gene families can result in an opening of phenotypic space in which previously cryptic mutations have strong effects, but also show how this expansion of phenotypic space begins to close as accumulated mutations in one part of a genetic network increasingly mask the effect of mutations in other parts.

## DISCUSSION

Here, we used natural and engineered cryptic cis-regulatory variation to identify and functionally dissect new transcription factor regulators of tomato inflorescence development in a four-gene regulatory network. Engineering additional cryptic cis-regulatory alleles with continuous epistatic effects and densely sampling thousands of inflorescences from hundreds of combinatorial genotypes allowed us to resolve the genetic architecture and genotype-phenotype map of this network. Mutations within this network tended to interact multiplicatively, but with even stronger positive synergistic (i.e. super-multiplicative) interactions within paralogous gene pairs, consistent with frequent redundancy between paralogs ^39,40^. Unexpectedly, we detected dose-dependent masking interactions acting simultaneously between paralog pairs, where mutations in one pair systematically shrink the effects of mutations in the other pair.

Pan-genome sequencing within and across taxa provides a rich resource for understanding how genetic networks are wired. Here, genetic variation in *S. habrochaites* and *S. pennellii* suggested the *PLT*s as candidate regulators of inflorescence morphology in tomato. However, pan-genome sequencing has also revealed extensive and diverse forms of variation whose molecular, developmental, and evolutionary significance remains unknown. Widespread structural variation in cis-regulatory regions and the frequent duplication and loss of both small and large genomic regions both drive quantitative expression variance and gene dosage ^18,41,42^. The dynamic emergence, divergence, and turnover of paralogous genes can alter the architectures and buffering of regulatory networks, thus perturbing component dosage and potentiating phenotypic change from canalized states when cryptic variants in paralogs converge.

Our detailed dissection of genetic architecture revealed how the *PLT-SEP* network can shift from a canalized state to one poised to release both subtle and substantial phenotypic change. We observed a hierarchical structure of genetic interactions: classical synergistic interactions within paralog pairs, driven by partial redundancy, combine via a multilinear interaction between the two paralog pairs. This interaction is further transformed by an exponential mapping that accounts for these mutations, which alter meristem maturation and development, combining multiplicatively to determine average branches per inflorescence. Our finding of a multilinear interaction between the two paralog pairs is notable because, while the multilinear model has been explored theoretically ^35,43,44^ and often is used to estimate how directional selection changes additive genetic variance ^45,46^, it has received limited empirical support for capturing real-world patterns of epistasis ^45,47^. Nonetheless, the multilinear model can be viewed as a continuous relaxation of Boolean (binary ‘on’ or ‘off’) gene regulatory models ^48^, accommodating a spectrum of allelic strengths and dosage effects rather than treating gene expression as a binary. We thus hypothesize that synergistic interactions within paralogous gene families, combined via multilinear interactions that reflect the structure and functional logic of gene regulatory networks, are likely a common genetic architecture that governs how phenotypic space is simultaneously expanded and constrained.

Notably, the *PLT-SEP* genetic network has accumulated extensive genetic variation both within tomato and between Solanaceae species. *J2* is a relatively recent duplication missing in many Solanaceae ^25,26^. While the *PLT* paralogs are broadly retained, their redundancy relationships have likely diverged and vary between genotypes and species. In contrast, in the Brassicaceae, while *PLT3* and *PLT7* orthologs are conserved, *J2* and *EJ2* MADS-box paralogs have been lost. Interestingly, Arabidopsis *plt* mutants do not have altered inflorescence architecture and branching is constrained across species in the family ^49^. The architecture of *PLT-SEP* regulatory networks in the Solanaceae may allow cryptic variation to accumulate more readily than in the Brassicaceae, priming epistasis and enhancing evolvability of branching. Testing this hypothesis will require broader genome sampling, uncovering patterns of paralog emergence, diversification, and loss, and identifying causal variation across species in these families. Critically, as we show (**Fig. 1 and 3**), this approach can test specific hypotheses while revealing additional components of conserved and diverged gene regulatory networks. More broadly, the principles uncovered here—where varying paralog redundancy relationships and presence-absence variation shape trait evolvability through genetic interactions that follow the multilinear model—likely extend across regulatory networks underlying other developmental and physiological programs, influencing the evolutionary trajectories of many traits.

Finally, a key aspect of our study was engineered genetic variation that densely sampled genotypic and phenotypic space in a controlled isogenic background. This approach provided the resolution needed to define the character and quantitative form of gene action and epistasis, represented as a surface illustrating how phenotypic effects from mutations within a paralog family combine when incorporating mutations across gene families (**Fig. 4g**). Detailed mapping of genetic interactions in this way could help reconcile the observation of widespread epistasis in model organisms with the challenge of detecting epistatic effects from allelic variation in natural populations ^50,51,12–15^. Placing natural alleles and their combinations onto similar surfaces could reveal how interactions among standing variants push populations into regions of genotypic space where phenotypic variation is either amplified, suppressed, or both. Beyond evolutionary insights, this framework has practical implications in crop engineering ^26,52,53^. Understanding how distinct genetic combinations, along with the specific forms of epistasis they engender, can yield the same phenotypic outcomes may inform targeted editing strategies to predictably ‘tune’ these interactions. By shifting populations or individuals to advantageous positions on the genotype– phenotype surface, such strategies could minimize undesirable pleiotropic effects and circumvent genetic constraints imposed by natural alleles in both complex breeding populations and elite genotypes. Realizing these opportunities will hinge on future research that encompasses larger networks of interacting genes, emphasizing how taxon-specific complements of paralogous genes and their variants shape network architecture—and hierarchical epistasis—across broader evolutionary clades.

## ONLINE CONTENT

## Reporting summary

Further information on research design is available in the Nature Portfolio Reporting Summary linked to this article.

## Data availability

All data are available within this Article and its Supplementary Information. RNA sequencing data is available at the Gene Expression Omnibus, GSE289537. The raw data with the number of branching events for each plant and inflorescence is provided in Supplementary Tables. All unique biological materials used in this manuscript are available for distribution upon request.

## Code availability

Code to reproduce the statistical analysis, quantitative modeling and the derived Figures is available at https://github.com/cmarti/tomato_branching/tree/master.

### Acknowledgements

We thank members of the Lippman laboratory for their support, enthusiasm, and helpful feedback, S. Hutton for support with field growing support, T. Mulligan, K. Schlecht, S. Qiao for assistance with plant care, and A. Le Rouzic for helpful conversations. SGZ is supported by the National Institutes of Health grant K99GM149939. YE is supported by the Israel Science Foundation grant 2390/23. CMG and DMM were supported by NIH grant R35 GM133613 and additional funding from the Simons Center for Quantitative Biology at Cold Spring Harbor Laboratory. ZBL and MB are supported by The National Science Foundation Plant Genome Research Program grant IOS-2129189, MB is supported by a Gatsby Charitable Foundation grant (GAT3731/GLK), and ZBL is supported by The National Science Foundation Plant Genome Research Program grant IOS-2216612 and the Howard Hughes Medical Institute.

## Author Contributions

Conceptualization: SGZ, DMM, ZBL

Formal analysis: SGZ, CM-G

Funding acquisition: SGZ, YQ, YE, DMM, ZBL

Investigation: SGZ, CM-G, BF, CPDC, ML, BS, AH, SS

Methodology: SGZ, CM-G, DMM, ZBL

Project administration: SGZ, CM-G, YE, DMM, ZBL

Writing – original draft: SGZ, CM-G, YE, DMM, ZBL

Writing – review & editing: SGZ, CM-G, BF, MB, YE, DMM, ZBL

## Additional information

The online version contains supplementary material available at https://doi.org/xxxxxx

## METHODS

### Plant materials

Seeds of wild type *Solanum lycopersicum* (cultivar M82, LA3475), *Solanum habrochaites* (LA1777), and *Solanum pennellii* (LA0716) were from our stocks. Introgression line IL3-4 (*Solanum pennellii* chromosome 3 introgressed into M82, LA4046) was obtained from the Tomato Genome Resource Center (Department of Plant Sciences, University of California at Davis) and the variant was validated by PCR amplification and Sanger sequencing (all primers used in this study are reported in **Supplementary Table 23**) ^54^. Two overlapping *Solanum habrochaites* chromosome 3 introgression lines LA3925 and LA3926, introgressed into tomato cultivar TA209, were obtained from the Tomato Genome Resource Center (Department of Plant Sciences, University of California at Davis) ^55^. Upon validation by PCR amplification and Sanger sequencing, only LA3925 contained the *ShEJ2^pro-3^*variant of interest, so LA3926 was used as a control for crosses between M82 and the introgressed region in the TA209 background. Mutants *j2-TE ej2^w^*and *j2 ej2* were from our stocks, as previously described ^24^.

### Motif enrichment and variant discovery

FIMO motif enrichment was performed on the sequence of open chromatin regions in the tomato meristem upstream of *SlEJ2* using the *Arabidopsis thaliana* non-redundant motif database curated at Plant TFDB (p value < 0.00001 and q-value < 0.01) ^56^. The same regions were used to search for insertion-deletion (indel) variants called previously from the tomato pangenome ^27^. Indels overlapping with annotated motifs were confirmed to not exist in linkage with previously reported *EJ2* variants (*ej2^w^, sb3*) by PCR and then used for subsequent experiments (Supplementary Table 1) ^24,57^.

### Genome editing

CRISPR-Cas9 mutagenesis and generation of transgenic tomato plants were performed following our standard protocol ^58^. Briefly, guide RNAs (gRNAs) were designed using the Geneious Prime software (https://www.geneious.com/) (gRNAs used in this study are listed in **Supplementary Table 23**). For Cas9 multiplex editing, the Golden Gate cloning system was used to assemble the binary vector containing the Cas9 and the specific gRNAs ^58,59^. For SpRY editing, vectors were constructed through a modular Gateway^TM^ assembly, as described previously (Invitrogen)^60^. Final binary vectors were then transformed into the tomato cultivar M82 by *Agrobacterium tumefaciens*-mediated transformation through tissue culture ^61^. First-generation transgenic plants (T0) were genotyped with specific primers surrounding the target sites (all primers used in this study are reported in **Supplementary Table 23**). To purify alleles from potential spontaneous mutations or CRISPR-Cas9 off-target effects following plant transformation, all T0 transgenic lines were backcrossed (BC1) to parental wild-type plants. BC1 populations were then screened by PCR and Kanamycin herbicide susceptibility for plants lacking the Cas9 transgene, PCR products of the targeted regions were Sanger sequenced to confirm inheritance of alleles, and allele-specific genotyping assays were designed for genotyping in subsequent generations. Selected BC1 plants were self-fertilized to generate F2 populations, and these segregating populations were used to validate the phenotypic effects of each allele by co-segregation. F2 or F3 homozygous mutant plants were then used for subsequent crossing and quantitative phenotypic analyses.

### Growth conditions and phenotyping

Seeds were directly sown in soil in 96-cell plastic flats and grown to 4-week-old seedlings in the greenhouse. Seedlings were then transplanted to 4L pots in the greenhouse for crossing and bulking purposes or directly to the fields at Cold Spring Harbor Laboratory, New York or at The University of Florida Gulf Coast Research and Education Center. Greenhouse conditions are long-day (16 h light, 26-28 °C / 8 h dark, 18-20 °C; 40-60% relative humidity) with natural light supplemented with artificial light from high-pressure sodium bulbs (∼250 μmol m^−2^ s^−1^). Plants in the fields were grown under drip irrigation and standard fertilizer regimes, and were used for quantifications of inflorescence branching, fruit shape, and sepal length.

To quantify inflorescence branching, inflorescences were counted in order of emergence in two rounds, approximately 60 days post sowing and 75 days post sowing. When available, four primary inflorescences and six axillary inflorescences were counted per plant. 60 or fewer branches were counted, if branching exceeded 60, “too many to count” (“TMTC”) was recorded and the number of branching events was treated as 60 for downstream analysis. Proliferated meristem in the place of inflorescence was indicated in the data as “proliferated”. Occasionally, inflorescences would fail to develop into countable structures, possibly due to stress, in which case, “inhibited” was recorded.

To quantify fruit shape, 10 fruits were collected at mature green stage, cut in transverse sections, and scanned on a single plane. The ratio of maximum height to width, Fruit Shape Index I, was determined from scanned images using Tomato Analyzer ^62^. To quantify sepal length, 10 closed mature floral buds of similar developmental stage (1-2 days before anthesis, i.e., before flower opening) per genotype were collected, length of sepals and petals were manually measured and the sepal/petal ratio was calculated ^24^.

### Phylogenetic Trees

Protein sequences of orthologs and paralogs of transcription factors (TF) of interest were obtained from Plant TFDB, MAFFT aligned, cleaned with BMGE, and a Maximum Likelihood Phylogeny was constructed with PhyML (NGPhylogeny.fr) ^63^. Trees were visualized with FigTree ^64^.

### RNA extraction and Illumina sequencing

Inflorescence meristems were collected from n = 4 plants at 8 weeks old under stereoscope magnification. Tissue was frozen, ground with beads, and RNA was extracted with TRIzol (Invitrogen) and a Direct-zol RNA Miniprep kit with on-column DNA digestion (Zymo Research). RNA was quantified with Qubit fluorimeter RNA HS assay kit (Invitrogen). Samples were treated with Ribo-Zero rRNA removal kit (Epicenter) and libraries prepared with a TruSeq V2 RNA-Seq prep kit (Illumina).

### RNA Sequencing Analysis

Published RNA-seq data of wild-type M82, *ej2*, *j2*, and *anantha*, mutant meristems were downloaded from SRA PRJNA376115, and PRJNA343677 ^21,34^. Reads were trimmed with Trimmomatic (ILLUMINACLIP:TruSeq2-PE.fa:2:30:10:1:FALSE LEADING:3 TRAILING:3 SLIDINGWINDOW:4:15 MINLEN:36) and aligned to the cDNA annotation of the reference genome sequence of tomato (SL4.0) using STAR v2.6.1.d ^65^. Normalization and quantification of individual transcript expression was done in R by calculating transcripts per million (TPM). Differential expression was calculated in R by DESeq2 and the top 200 most differentially expressed genes across wild type meristem maturation were used for principal component analysis of all meristem samples using Python scikit-learn PCA.transform ^66^.

### Dual Luciferase assay

A Gateway^TM^-compatible dual-luciferase reporter vector (pSZ106) was assembled using the MoClo GoldenGate assembly system ^59,67^. Briefly, a Gateway^TM^ AttR4-AttL1R cassette (Invitrogen) was cloned upstream of a 46 base pair minimal 35S promoter driving the Firefly luciferase coding sequence (pICSL80001-pL0_fLUC-I (CDS1)) with a Nopaline Synthetase terminator (pICH41421) ^59,67^. A Cauliflower Mosaic Virus 35S promoter (pICH51266) was cloned upstream of the coding sequence of Renilla luciferase (pSB123 - pL0_rLUC-I (CDS1), Addgene) with a Nopaline Synthetase terminator (pICH41421) ^67^. Both luciferase expression cassettes were cloned into the pICSL4723 binary vector backbone with an NPTII selection cassette. *SlEJ2^pro-3^*, *ShEJ2^pro-3^* and *SpEJ2^pro-3^* alleles were cloned into pDONR™ P4-P1r and introduced into pSZ106 by Gateway^TM^ cloning (Invitrogen). *SlERF12* (Solyc02g077840), *SlPLT380* (Solyc05g051380), and *SlPLT710-short* (Solyc11g010710) were cloned into pDONR207 and introduced into pEAQ-HT-DEST3 by Gateway^TM^ cloning (Invitrogen).

All binary expression vectors were transformed into *Agrobacterium tumefaciens* and cultured at 28°C overnight in selective media. Overnight cultures were diluted and grown at 28°C to OD600 1 AU, spun down, and washed into inductive media (10mM MES pH 5.7, 10mM MgCl_2_, 100µM 3’,5’-Dimethyoxy-4’-hydroxyacetophenone) at OD600 1 AU. Bacteria was induced for 3 hours lying horizontally at the bench, then equal volumes of promoter and TF media were combined and co-infiltrated into young fully expanded leaves of 4 week old *Nicotiana benthamiana* plants grown in long days (16 h light / 8 h dark, 22°C; 40-60% relative humidity). Plants were returned to the growth chamber and 100mg tissue was collected and frozen for measurement 3 days after infiltration.

Luciferase activity was measured using a Dual Luciferase Reporter Assay System kit (Promega) according to Moyle *et al.* ^68^. Briefly, tissue was homogenized in a Spex Sample Prep 2010 Geno/Grinder (Cole Parmer) and 10 mg of tissue powder was mixed with 100 µl of passive lysis buffer (Promega). Cellular debris was pelleted at 7,500 × g for 1 min, and supernatant was diluted 40X in passive lysis buffer and 15 µl of sample was transferred to 3 replicate wells of a white flat-bottom Costar 96-well plate (Corning). The assay was measured using a GloMax 96 microplate luminometer, and 75 µl per well of luciferase assay reagent and Stop & Glo reagent were added and measured stepwise (Promega).

### Segregating Populations

BC1 inbred plants of genotype *EJ2pro j2* were crossed to BC1 inbred plants of genotype *plt3 plt7/+* and genotyped in the F1 generation by allele specific PCR to determine the presence of all desired alleles. F2 seed was sown and tissue was collected for DNA extraction and transplanted without genotyping. Genotype of plants was confirmed after phenotyping by allele-specific PCR.

### Linear regression models

Phenotypic data was summarized at the plant level for quantitative modeling, with abnormal inflorescences marked as “proliferated” or “inhibited” excluded from further analysis. For each plant *i*, we consider the total number of branching events across all inflorescences *y_i_*, and the number of inflorescences *t_i_*. The total number of branching events *y_i_* was modeled as being either Poisson or negative binomially distributed with exposure *t_i_*. We defined a basis for the constant component, defined by a vector that takes the value 1 for each plant, and a basis for the additive model by expanding this constant basis to also include an additive and dominant component for each locus. The additive component was coded as –1, 0, 1 for homozygous wild-type, heterozygous mutant and homozygous mutant, respectively; whereas the dominant component was coded as 1 for heterozygous genotypes and 0 elsewhere. A basis for pairwise interaction models was built by extending the basis for the additive model with additional basis vectors composed by taking the product of each possible pair of additive and dominance components ^69^. Models were defined and fit using *statsmodels* ^70^ python package using Poisson and negative binomial likelihoods with the identity and log link functions. Genotype-season Maximum Likelihood Estimates (MLE) for the number of branching events were obtained by defining a dummy variable for each genotype-season combination that took a value of 1 for plants of that genotype and 0 otherwise, while assuming that all genotypes share the overdispersion parameter for the negative binomial likelihood function representing plant-to-plant variability that is jointly estimated with the genotype-season means. Confidence intervals for the MLEs were derived using *statsmodels* with a log-link between model parameters (genotype estimates) and the average number of branching events.

### Hierarchical model

Data was modeled with a negative binomial likelihood function as explained in the *Linear regression models* section. However, for the hierarchical model, the log-transformed genotype means log(μ) are given by a multilinear function log(μ) = θ_WT_ + φ_PLT_ + φ_SEP_ - θ_Int_ φ_PLT_φ_SEP_ where θ_WT_ is the wild-type log-transformed expected branching events, φ_PLT_ controls the log effect of the relevant combination of mutations in *PLT3* and *PLT7* when placed in a wild-type *EJ2 J2* background, φ_SEP_ controls the log effect of the relevant combination of mutations in *EJ2* and *J2* combinations in a wild-type *PLT3 PLT7* background and θ_Int_ represents the masking interaction between the two phenotypes ^35^. Note that φ_PLT_ takes a different values for every possible combination of mutations in *PLT3* and *PLT7*, while φ_SEP_ takes a different value for every possible combination of mutations in *EJ2* and *J2*, so that inferring the values for φ_PLT_ and φ_SEP_ is equivalent to allowing a full set of additive, dominance and pairwise interactions within each of *PLT3/PLT7* and *EJ2*/*J2*. Thus, the hierarchical model is equivalent to fitting pairwise interactions within paralog pairs, then combining the within pair effects via a multilinear interaction across pairs, and then transforming the result via an exponential function. This model was coded in PyTorch ^71^ and the Maximum Likelihood solution was found running the Adam optimizer for 10,000 iterations and checking for convergence.

## EXTENDED DATA FIGURE

**Extended Data Figure 1:**
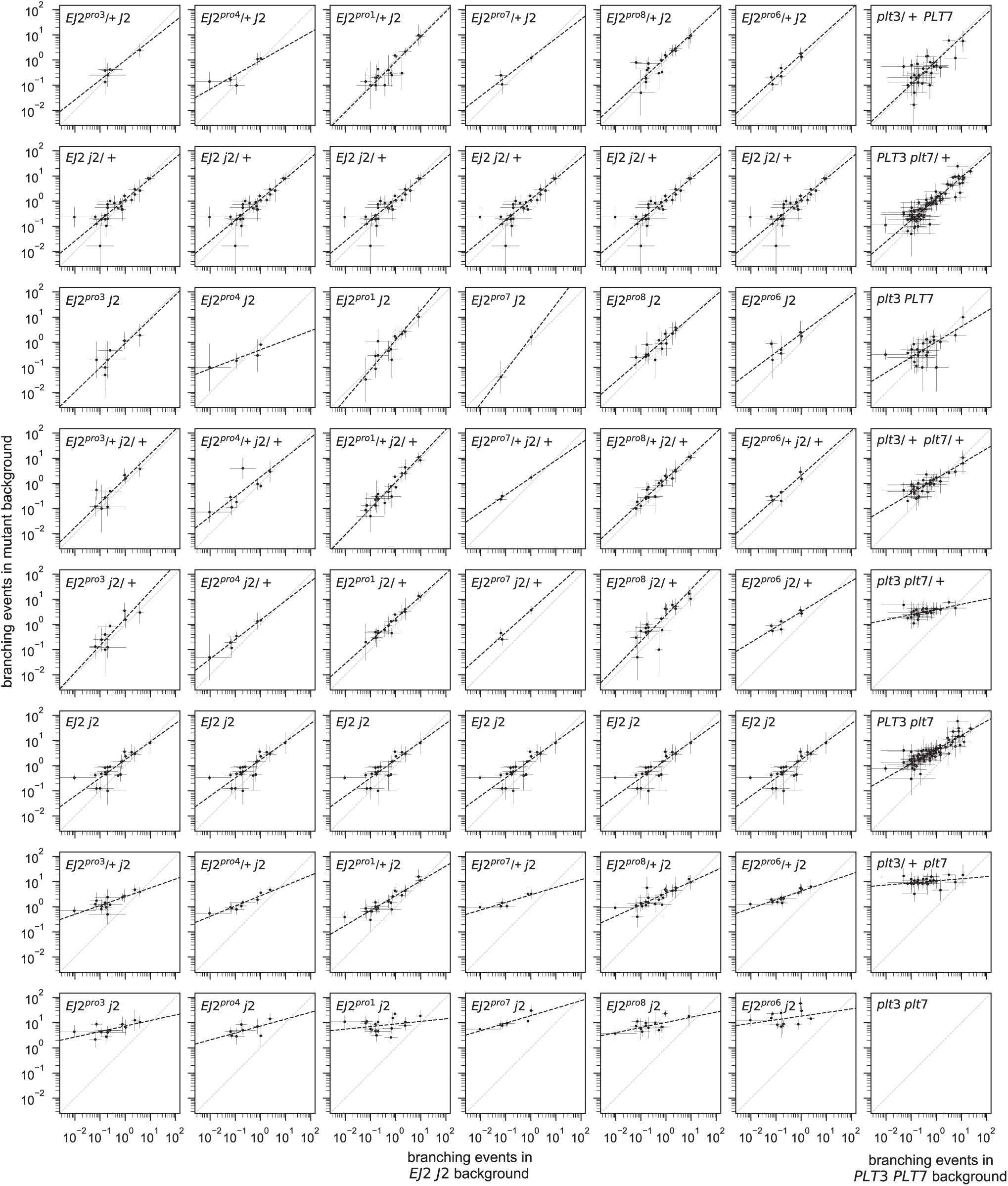
Mutations in one paralog pair linearly re-scale the effects of mutations in the other paralog pair. Scatterplots representing the Maximum Likelihood Estimate (MLE) of the expected number of branching events. Every dot represents a possible combination of mutations in *PLT3* and *PLT7* (first 6 columns) or *EJ2* and *J2* (last column) in a specific season. The value plotted on the y-axis corresponds to the phenotype conferred by the given combinations of mutations either in the designated background at *EJ2* and *J2* (first 6 columns) or the designated background at *PLT3* and *PLT7* (last column); x-axis values are given by the phenotype of each set of mutations in the wild-type background. Error bars represent the 95% confidence interval for the MLEs. Total least squares regression lines for the log MLEs are represented with black dashed lines. Genotype-season combinations with a 95% confidence interval wider than a thousand-fold range are not shown.

**Extended Data Figure 2:**
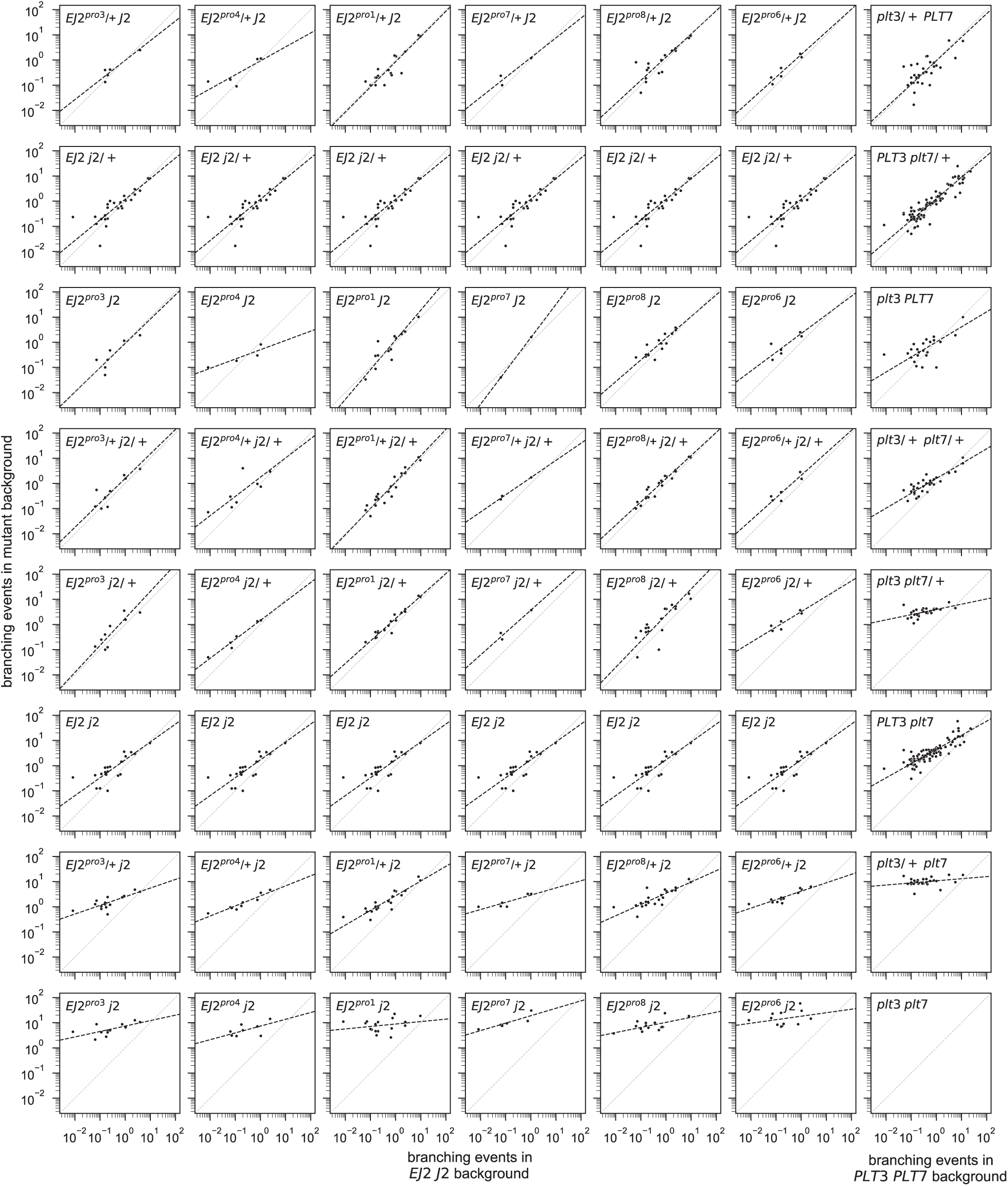
Linear rescaling of mutational effects observed between paralog pairs is robust under an alternative method for estimating genotype-season means. Scatterplots representing the sample average number of branching events. Every dot represents a possible combination of mutations in *PLT3* and *PLT7* (first 6 columns) or *EJ2* and *J2* (last column) in a specific season. The value plotted on the y-axis corresponds to the sample average phenotype conferred by the given combinations of mutations either in the designated background at *EJ2* and *J2* (first 6 columns) or the designated background at *PLT3* and *PLT7* (last column); x-axis values are given by the sample average of each set of mutations in the wild-type background. Total least squares regression lines for the log sample means are represented with black dashed lines. Genotype-season combinations in which no branching was observed are not shown.

**Extended Data Figure 3:**
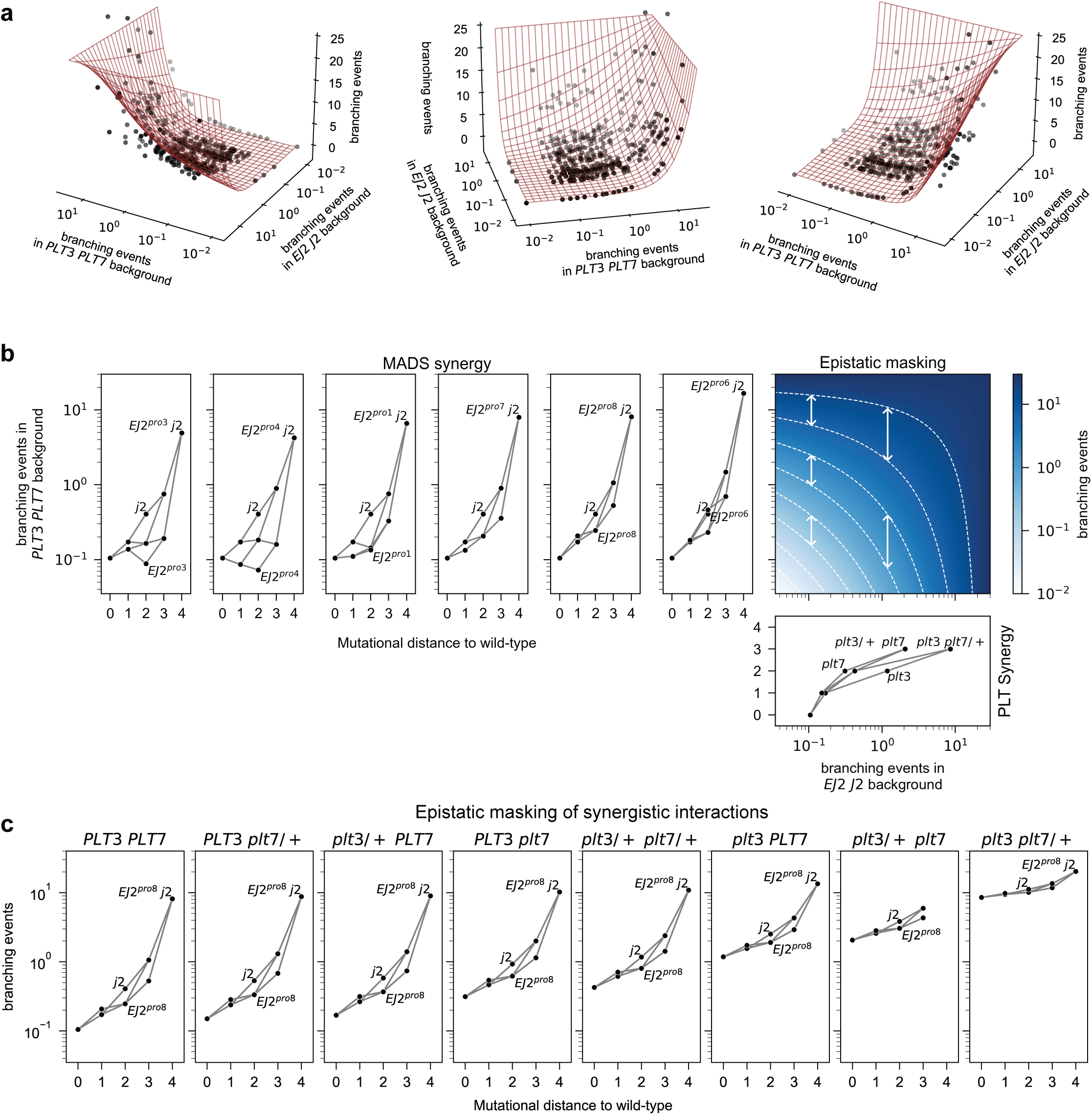
Understanding synergy and masking under the inferred hierarchical model. **(a)** Different views of the two-dimensional surface representing the expected number of branching events as a function of the mean branching events conferred by placing the combination of mutations in each paralog pair in the wild-type background of the other pair. Points represent the independent maximum likelihood estimates (MLEs) for the expected number of branching events for the measured genotype-season combinations. **(b)** Complete representation of the inferred hierarchical model of gene interaction. Left and bottom panels represent the estimated number of branching events for each paralog pair combination in the wild-type background of the other pair as a function of the Hamming distance to the wild-type genotype. Different *EJ2^pro^* alleles show phenotypic effects of different magnitude but all interact with *J2* in a similar way. The phenotype for any genotype is obtained by combining the independent effect of the mutations in each pair of paralogs through the two-dimensional surface in the middle panel. This heatmap represents the inferred multilinear function that quantitatively characterizes epistatic masking between PLT and MADS genes. White dashed lines represent isophenotypic lines, this is, combinations of background phenotypic effects that result in the same number of branching events when combined. Note that the function is linear across any horizontal or vertical transect, as illustrated by the constant distance between isophenotypic lines across any transect. Deviation from a slope of -1 in the shape of the isophenotypic lines, which are scale-independent, indicates the presence of an epistatic interaction between the two pairs of paralogs. Note that distance between isophenotypic lines increases in highly mutated backgrounds, indicating that a larger perturbation is required to achieve the same phenotypic outcome. **(c)** Estimated number of branching events for all *EJ2^pro8^ J2* combinations across genetic backgrounds with an increasing number of mutations in *PLT3* and *PLT7* illustrates how the synergistic interactions between *EJ2^pro8^* and *j2* become masked as *PLT3* and *PLT7* become increasingly mutated.

## SUPPLEMENTARY DATA FIGURE

**Supplementary Figure 1:**
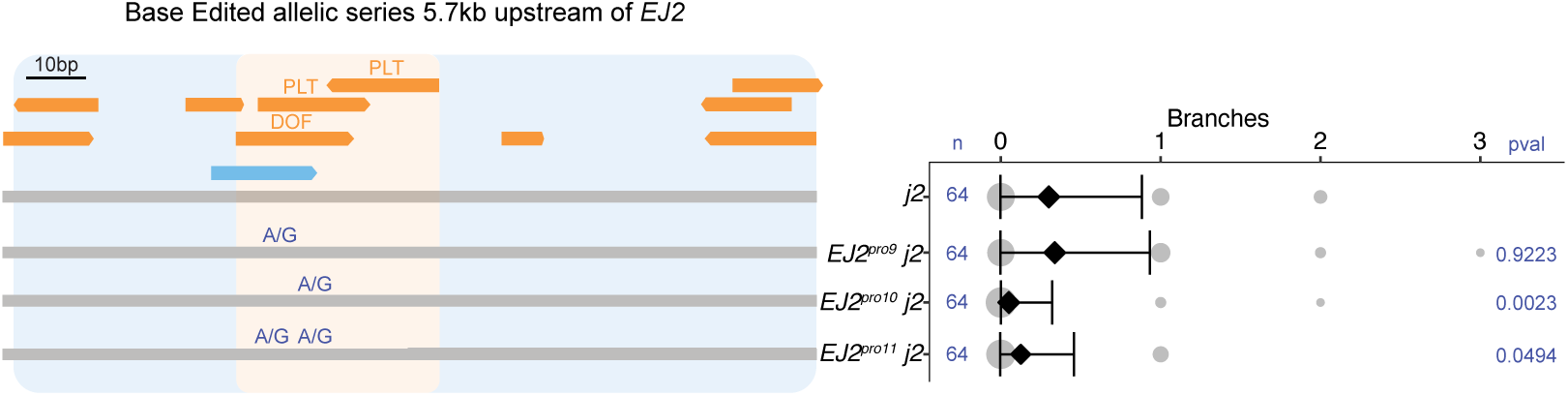
Base edited *EJ2 cis*-regulatory alleles. Left: 153 bp target region located 5.7kb upstream of the *EJ2* transcription start site showing annotated TFBSs (orange), including the focal DOF and AP2/ERF binding sites. Also shown is a CRISPR/dCas9-ABE8E base editor gRNA (light blue) used for targeted mutagenesis to generate an allelic series (*EJ2^pro^*). SNVs, blue. Right: Quantifications of inflorescence branching for each genotype. Area of grey circles, numbers of inflorescences; black diamonds and bars, mean values and standard deviations. Total number (n) of observations and adjusted p-values from Two-sided Dunnett’s Compare with Control Test.

**Supplementary Figure 2:**
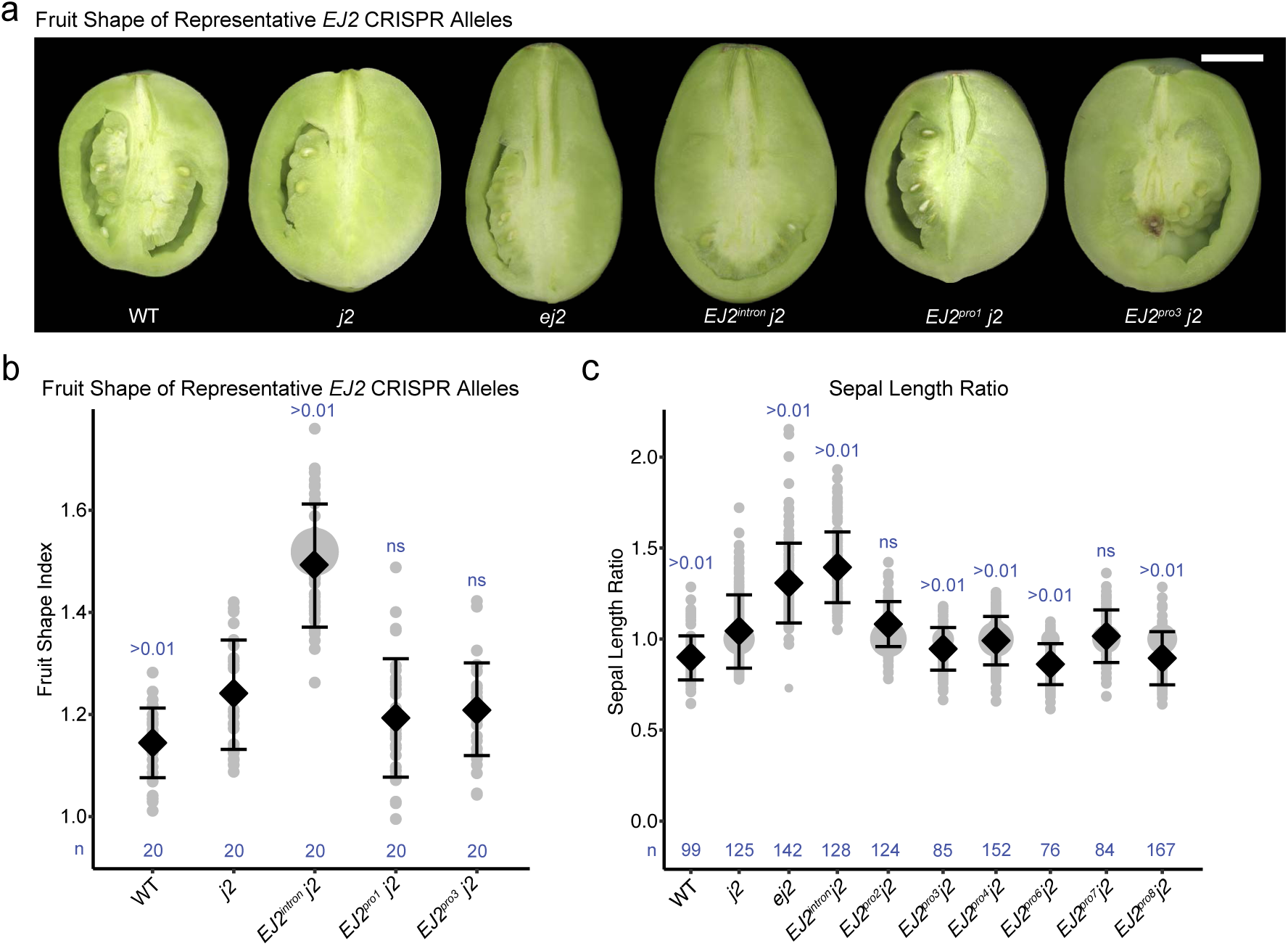
Fruit shape and sepal length phenotypes of *EJ2 cis*-regulatory alleles. **(a)** Transverse sections of representative mature green fruit. Scale bar, 1 cm. **(b)** Ratio of maximum height to width of mature green fruits (Fruit Shape Index I)^62^. Grey circles, index, scaled by number of fruits; black diamonds and bars, mean values and standard deviations. Total number (n) of fruits and adjusted p-values from Two-sided Dunnett’s Compare with Control Test. **(c)** Ratio of sepal length allele/*j2* background. Grey circles, ratio, scaled by number of flowers; black diamonds and bars, mean values and standard deviations. Total number (n) of flowers and adjusted p-values from Two-sided Dunnett’s Compare with Control Test.

**Supplementary Figure. 3:**
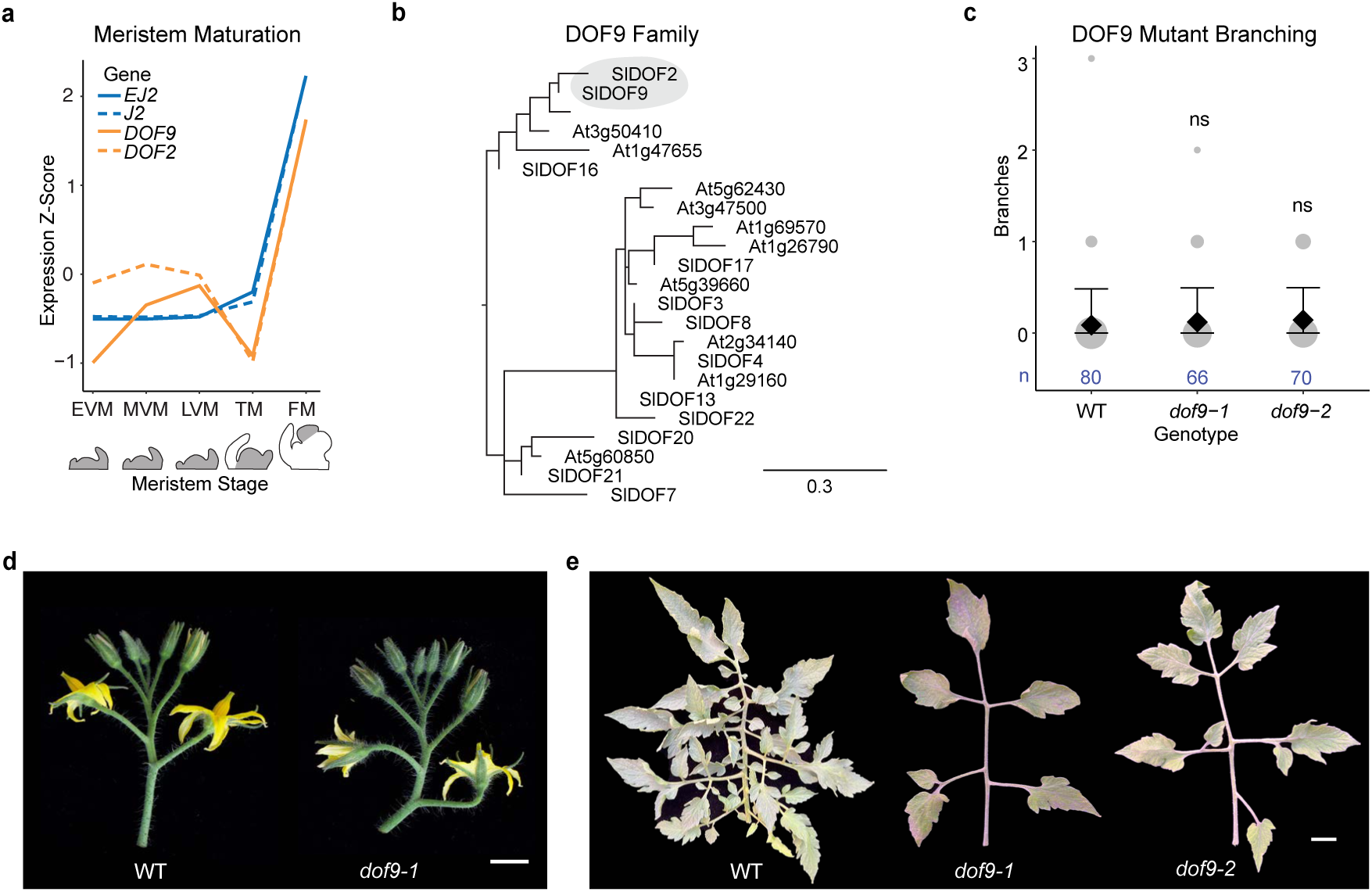
*DOF9* transcription factor mutant has no phenotype in inflorescence architecture. **(a)** Meristem expression dynamics of *EJ2, J2, DOF9,* and *DOF2* over five stages (depicted in grey): Early Vegetative Meristem (EVM), Middle (MVM), Late (LVM), Transition (TM), and Floral Meristem (FM). Despite similar expression dynamics, *DOF2* is lowly expressed (9.06 transcripts per million (TPM) in LVM, lower than associated LVM leaf primordia samples at 25.25 TPM) ^34^. **(b)** Phylogenetic tree of DOF9 ortholog proteins from Arabidopsis (*At*) and tomato (*Sl*). DOF9 and its paralog DOF2 are circled. Scale bar is relative substitution rate. **(c)** Quantification of branching in *dof9* mutants. Area of grey circles correspond to the number of inflorescences, black diamonds are mean values and bars are standard deviations. Number (n) of observations and p-values from Two-sided Dunnett’s Compare with Control Test shown in blue. **(d)** Representative inflorescences of WT and *dof9* mutant. Scale bar is 1 cm. **(e)** Representative fifth leaves of WT and *dof9* mutants. Scale bar is 1 cm.

**Supplementary Figure 4:**
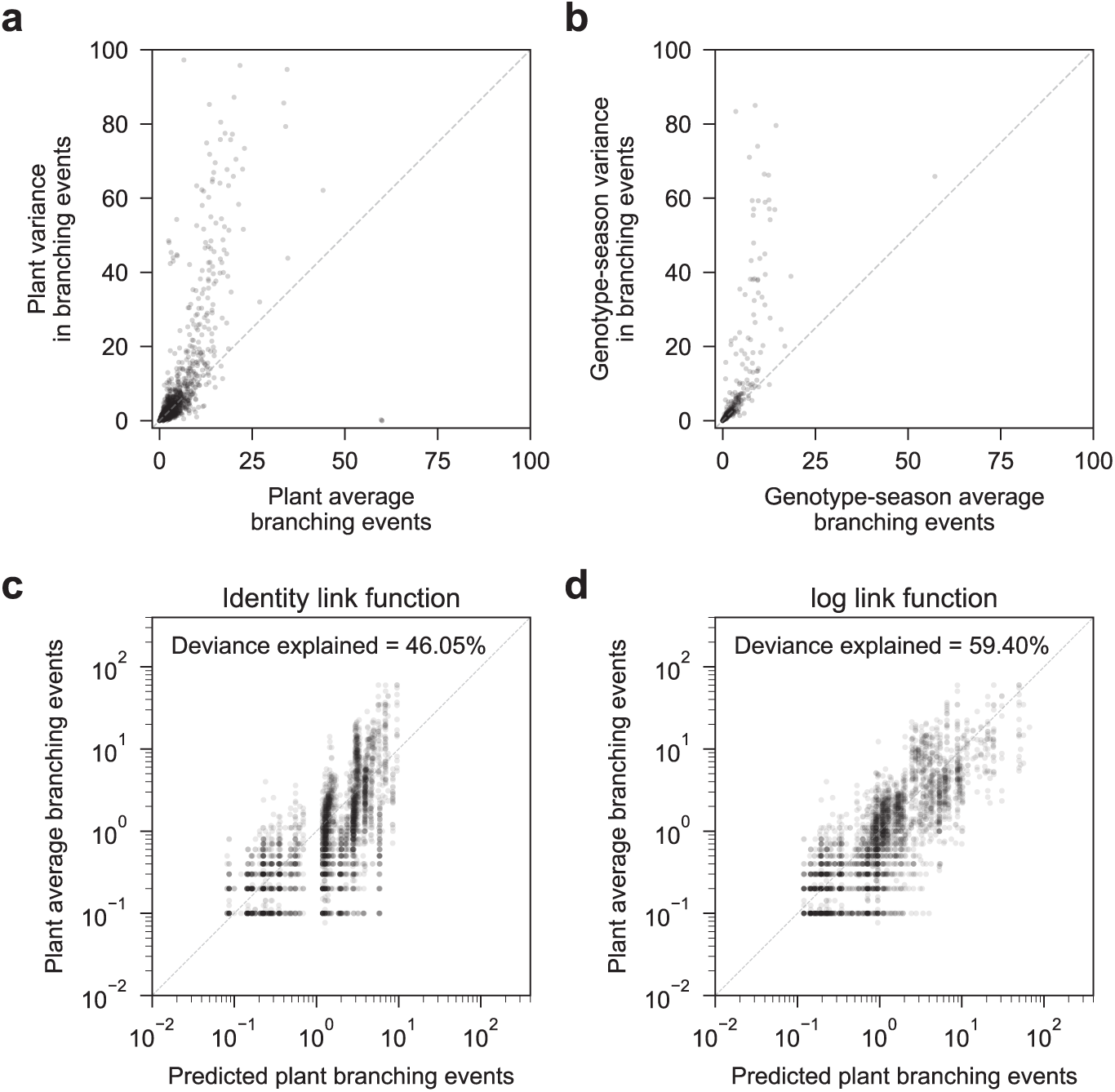
Models for the number of branching events. **(a)** Scatterplot of the per-plant average number of branching events and the variance across inflorescences within the same plant. Dashed line shows expectation under the assumption that branching events are Poisson distributed. **(b)** Scatterplot of the per-genotype and season average number of branching events and the variance across inflorescences within the same genotype and season. Dashed line shows expectation under the assumption that branching events are Poisson distributed. **(c)** Scatterplot showing the predicted branching events under an additive Poisson regression model with identity link function and observed per-plant average number of branching events. **(d)** Scatterplot showing the predicted branching events under an additive Poisson regression model with log link function (i.e. assuming that mutations combine multiplicatively) and observed per-plant average number of branching events.

**Supplementary Figure 5:**
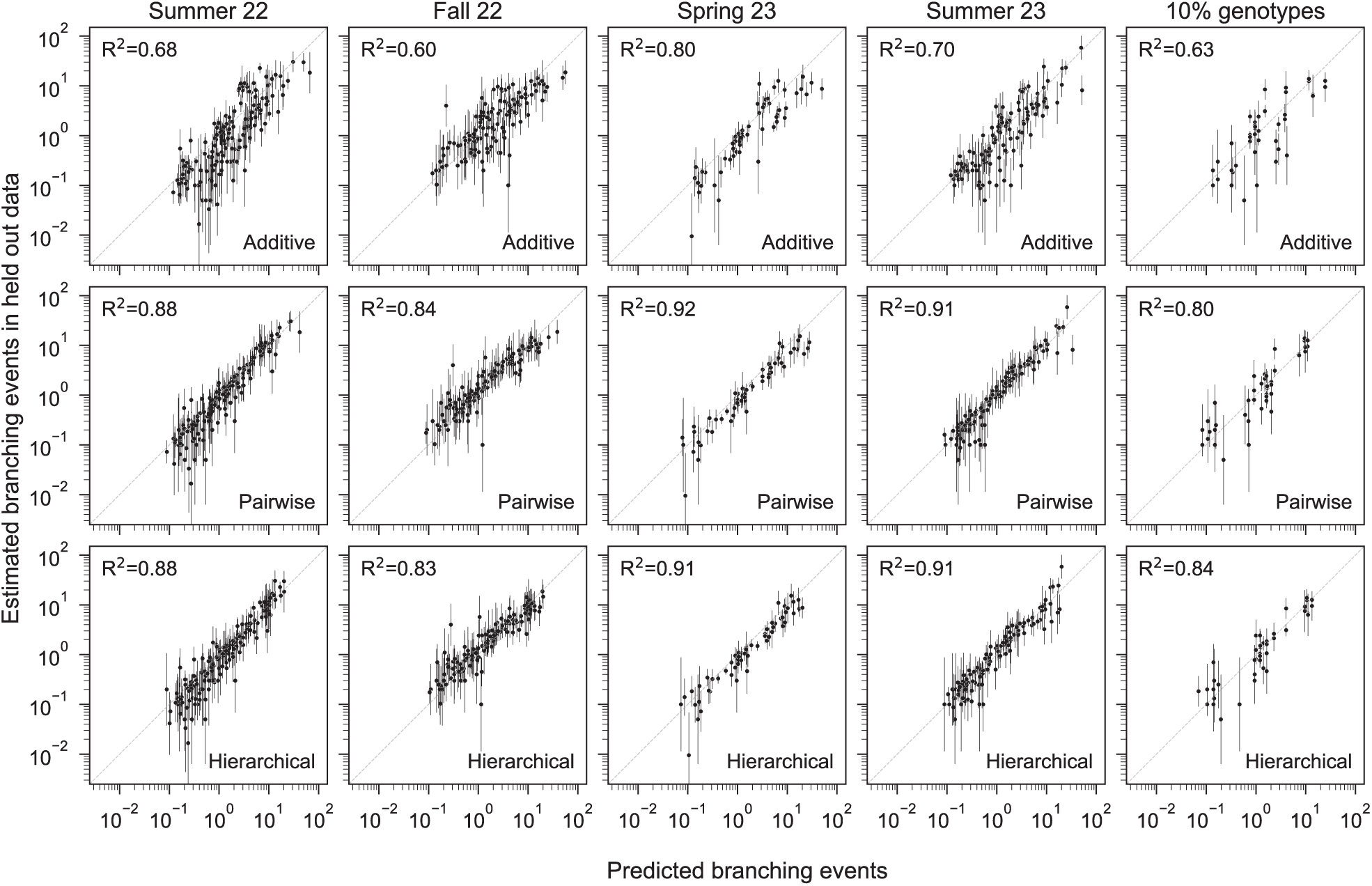
Predictive performance of additive, pairwise and hierarchical model on held-out seasons and genotypes. Scatterplots comparing the log expected number of branching events under different models with the log Maximum Likelihood Estimate (MLE) of the genotype-season means for different subsets of held-data. In each of the first 4 columns data from a single season is held-out, whereas in the last column the same random 10% of genotypes are held-out across all seasons. Error bars represent the 95% confidence interval for the MLEs. Genotype-season combinations with a 95% confidence interval wider than a thousand-fold range are not shown. The reported R^2^ values correspond to the squared Pearson coefficients between the log predicted number of branches and the log MLEs for the held-out data.

## REFERENCES

1. Waddington, C. H. Canalization of development and genetic assimilation of acquired characters. Nature 183, 1654–1655 (1959).

2. Rockman, M. V. The QTN program and the alleles that matter for evolution: all that’s gold does not glitter. Evolution 66, 1–17 (2012).

3. Lee, Y. W., Gould, B. A. & Stinchcombe, J. R. Identifying the genes underlying quantitative traits: a rationale for the QTN programme. AoB Plants 6, (2014).

4. Raju, A., Xue, B. & Leibler, S. A theoretical perspective on Waddington’s genetic assimilation experiments. Proc Natl Acad Sci U S A 120, e2309760120 (2023).

5. Gibson, G. & Dworkin, I. Uncovering cryptic genetic variation. Nat Rev Genet 5, 681–690 (2004).

6. Paaby, A. B. & Rockman, M. V. Cryptic genetic variation: evolution’s hidden substrate. Nat Rev Genet 15, 247–258 (2014).

7. Mackay, T. F. C. & Anholt, R. R. H. Pleiotropy, epistasis and the genetic architecture of quantitative traits. Nat Rev Genet 25, 639–657 (2024).

8. Frank, S. A. Robustness and complexity. Cell Syst 14, 1015–1020 (2023).

9. Schneider, R. F. & Meyer, A. How plasticity, genetic assimilation and cryptic genetic variation may contribute to adaptive radiations. Mol Ecol 26, 330–350 (2017).

10. Ehrenreich, I. M. & Pfennig, D. W. Genetic assimilation: a review of its potential proximate causes and evolutionary consequences. Ann Bot 117, 769–779 (2016).

11. Sangster, T. A. et al. HSP90 affects the expression of genetic variation and developmental stability in quantitative traits. Proc Natl Acad Sci U S A 105, 2963–2968 (2008).

12. Nadeau, J. H. Modifier genes in mice and humans. Nat Rev Genet 2, 165–174 (2001).

13. Carlborg, O. & Haley, C. S. Epistasis: too often neglected in complex trait studies? Nat Rev Genet 5, 618–625 (2004).

14. Mackay, T. F. C. Epistasis and quantitative traits: using model organisms to study gene-gene interactions. Nat Rev Genet 15, 22–33 (2014).

15. Sackton, T. B. & Hartl, D. L. Genotypic Context and Epistasis in Individuals and Populations. Cell 166, 279–287 (2016).

16. Wolter, F. & Puchta, H. Application of CRISPR/Cas to Understand Cis- and Trans-Regulatory Elements in Plants. Methods Mol Biol 1830, 23–40 (2018).

17. Liang, Z. & Schnable, J. C. Functional Divergence between Subgenomes and Gene Pairs after Whole Genome Duplications. Mol Plant 11, 388–397 (2018).

18. Birchler, J. A. & Yang, H. The multiple fates of gene duplications: Deletion, hypofunctionalization, subfunctionalization, neofunctionalization, dosage balance constraints, and neutral variation. Plant Cell 34, 2466–2474 (2022).

19. Mao, Y. et al. Structurally divergent and recurrently mutated regions of primate genomes. Cell 187, 1547–1562.e13 (2024).

20. Jeong, H. et al. Structural polymorphism and diversity of human segmental duplications. Nat Genet 57, 390–401 (2025).

21. Lemmon, Z. H. et al. The evolution of inflorescence diversity in the nightshades and heterochrony during meristem maturation. Genome Res 26, 1676–1686 (2016).

22. Hilgenhof, R. et al. Morphological trait evolution in Solanum (Solanaceae): Evolutionary lability of key taxonomic characters. TAXON 72, 811–847 (2023).

23. Périlleux, C. & Huerga-Fernández, S. Reflections on the Triptych of Meristems That Build Flowering Branches in Tomato. Front Plant Sci 13, 798502 (2022).

24. Soyk, S. et al. Bypassing Negative Epistasis on Yield in Tomato Imposed by a Domestication Gene. Cell 169, 1142–1155.e12 (2017).

25. Morel, P. et al. Divergent Functional Diversification Patterns in the SEP/AGL6/AP1 MADS-Box Transcription Factor Superclade. Plant Cell 31, 3033–3056 (2019).

26. Benoit, M. et al. Solanum pan-genomics and pan-genetics reveal paralogs as contingencies in crop engineering. bioRxiv 2024.09.10.612244 (2024) doi:10.1101/2024.09.10.612244.

27. Alonge, M. et al. Major Impacts of Widespread Structural Variation on Gene Expression and Crop Improvement in Tomato. Cell 182, 145–161.e23 (2020).

28. Zhou, Y. et al. Graph pangenome captures missing heritability and empowers tomato breeding. Nature 606, 527–534 (2022).

29. Hendelman, A. et al. Conserved pleiotropy of an ancient plant homeobox gene uncovered by cis-regulatory dissection. Cell 184, 1724–1739.e16 (2021).

30. Walton, R. T., Christie, K. A., Whittaker, M. N. & Kleinstiver, B. P. Unconstrained genome targeting with near-PAMless engineered CRISPR-Cas9 variants. Science 368, 290–296 (2020).

31. Ren, Q. et al. PAM-less plant genome editing using a CRISPR-SpRY toolbox. Nat Plants 7, 25–33 (2021).

32. Yamaguchi, N., Jeong, C. W., Nole-Wilson, S., Krizek, B. A. & Wagner, D. AINTEGUMENTA and AINTEGUMENTA-LIKE6/PLETHORA3 Induce LEAFY Expression in Response to Auxin to Promote the Onset of Flower Formation in Arabidopsis. Plant Physiol 170, 283–293 (2016).

33. Hu, G. et al. The auxin-responsive transcription factor SlDOF9 regulates inflorescence and flower development in tomato. Nat Plants 8, 419–433 (2022).

34. Park, S. J., Jiang, K., Schatz, M. C. & Lippman, Z. B. Rate of meristem maturation determines inflorescence architecture in tomato. Proc Natl Acad Sci U S A 109, 639–644 (2012).

35. Hansen, T. F. & Wagner, G. P. Modeling genetic architecture: a multilinear theory of gene interaction. Theor Popul Biol 59, 61–86 (2001).

36. Akaike, H. A new look at the statistical model identification. IEEE Trans. Automat. Contr. 19, 716–723 (1974).

37. Kenneth P. Burnham, D. R. A. Model Selection and Multimodel Inference. (Springer New York, 2010).

38. Domingo, J., Baeza-Centurion, P. & Lehner, B. The causes and consequences of genetic interactions (epistasis). Annu. Rev. Genomics Hum. Genet. 20, 433–460 (2019).

39. Diss, G., Ascencio, D., DeLuna, A. & Landry, C. R. Molecular mechanisms of paralogous compensation and the robustness of cellular networks. J Exp Zool B Mol Dev Evol 322, 488–499 (2014).

40. Iohannes, S. D. & Jackson, D. Tackling redundancy: genetic mechanisms underlying paralog compensation in plants. New Phytologist 240, 1381–1389 (2023).

41. Computational Pan-Genomics Consortium. Computational pan-genomics: status, promises and challenges. Brief Bioinform 19, 118–135 (2018).

42. Birchler, J. A. & Veitia, R. A. One Hundred Years of Gene Balance: How Stoichiometric Issues Affect Gene Expression, Genome Evolution, and Quantitative Traits. Cytogenet Genome Res 161, 529–550 (2021).

43. The role of epistatic gene interactions in the response to selection and the evolution of evolvability. Theoretical Population Biology 68, 179–196 (2005).

44. Le Rouzic, A., Álvarez-Castro, J. M. & Hansen, T. F. The evolution of canalization and evolvability in stable and fluctuating environments. Evol. Biol. 40, 317–340 (2013).

45. Le Rouzic, A. Estimating directional epistasis. Front Genet 5, 198 (2014).

46. Bourg, S., Bolstad, G. H., Griffin, D. V., Pélabon, C. & Hansen, T. F. Directional epistasis is common in morphological divergence. Evolution 78, 934–950 (2024).

47. Pavlicev, M., Le Rouzic, A., Cheverud, J. M., Wagner, G. P. & Hansen, T. F. Directionality of epistasis in a murine intercross population. Genetics 185, 1489–1505 (2010).

48. Evolvability and robustness: A paradox restored. Journal of Theoretical Biology 430, 78–85 (2017).

49. Mudunkothge, J. S. & Krizek, B. A. Three Arabidopsis AIL/PLT genes act in combination to regulate shoot apical meristem function. Plant J 71, 108–121 (2012).

50. Omholt, S. W., Plahte, E., Oyehaug, L. & Xiang, K. Gene regulatory networks generating the phenomena of additivity, dominance and epistasis. Genetics 155, 969–980 (2000).

51. Gjuvsland, A. B., Hayes, B. J., Omholt, S. W. & Carlborg, O. Statistical epistasis is a generic feature of gene regulatory networks. Genetics 175, 411–420 (2007).

52. Mascher, M., Jayakodi, M., Shim, H. & Stein, N. Promises and challenges of crop translational genomics. Nature 636, 585–593 (2024).

53. Schreiber, M., Jayakodi, M., Stein, N. & Mascher, M. Plant pangenomes for crop improvement, biodiversity and evolution. Nat Rev Genet 25, 563–577 (2024).

## References (Methods)

54. Alseekh, S. et al. Resolution by recombination: breaking up Solanum pennellii introgressions. Trends Plant Sci 18, 536–538 (2013).

55. Monforte, A. J. & Tanksley, S. D. Development of a set of near isogenic and backcross recombinant inbred lines containing most of the Lycopersicon hirsutum genome in a L. esculentum genetic background: a tool for gene mapping and gene discovery. Genome 43, 803–813 (2000).

56. Jin, J. et al. PlantTFDB 4.0: toward a central hub for transcription factors and regulatory interactions in plants. Nucleic Acids Res 45, D1040–D1045 (2017).

57. Soyk, S. et al. Duplication of a domestication locus neutralized a cryptic variant that caused a breeding barrier in tomato. Nat Plants 5, 471–479 (2019).

58. Rodríguez-Leal, D., Lemmon, Z. H., Man, J., Bartlett, M. E. & Lippman, Z. B. Engineering Quantitative Trait Variation for Crop Improvement by Genome Editing. Cell 171, 470– 480.e8 (2017).

59. Engler, C. et al. A Golden Gate Modular Cloning Toolbox for Plants. (2014) doi:10.1021/sb4001504.

60. Lowder, L. G. et al. A CRISPR/Cas9 Toolbox for Multiplexed Plant Genome Editing and Transcriptional Regulation. Plant physiology 169, (2015).

61. Van Eck, J., Kirk, D. D. & Walmsley, A. M. Tomato (Lycopersicum esculentum). Agrobacterium Protocols 459–474 (2006).

62. Brewer, M. T. et al. Development of a controlled vocabulary and software application to analyze fruit shape variation in tomato and other plant species. Plant Physiol. 141, 15–25 (2006).

63. Lemoine, F., et al. NGPhylogeny.fr: new generation phylogenetic services for non-specialists. Nucleic acids research 47, (2019).

64. FigTree. http://tree.bio.ed.ac.uk/software/figtree/.

65. Dobin, A. et al. STAR: ultrafast universal RNA-seq aligner. Bioinformatics (Oxford, England) 29, (2013).

66. Pedregosa, F. et al. Scikit-learn: Machine Learning in Python. Journal of Machine Learning Research 12, 2825–2830 (2011).

67. Mortensen, S. et al. EASI Transformation: An Efficient Transient Expression Method for Analyzing Gene Function in Seedlings. Front Plant Sci 10, 755 (2019).

68. Moyle, R. L. et al. An Optimized Transient Dual Luciferase Assay for Quantifying MicroRNA Directed Repression of Targeted Sequences. Front Plant Sci 8, 1631 (2017).

69. Tiwari, H. K. & Elston, R. C. Deriving components of genetic variance for multilocus models. Genetic Epidemiology 14, 1131–1136 (1997).

70. Seabold, S. & Perktold, J. Statsmodels: Econometric and Statistical Modeling with Python. scipy (2010) doi:10.25080/Majora-92bf1922-011.

71. Paszke, A., et al. PyTorch: An Imperative Style, High-Performance Deep Learning Library. (2019).

